# Eddy current-induced artifacts correction in high gradient strength diffusion MRI with dynamic field monitoring: demonstration in ex vivo human brain imaging

**DOI:** 10.1101/2023.02.15.528684

**Authors:** Gabriel Ramos-Llordén, Daniel Park, John E. Kirsch, Alina Scholz, Boris Keil, Chiara Maffei, Hong-Hsi Lee, Berkin Bilgiç, Brian L. Edlow, Choukri Mekkaoui, Anastasia Yendiki, Thomas Witzel, Susie Y. Huang

## Abstract

**Purpose:** To demonstrate the advantages of spatiotemporal magnetic field monitoring to correct eddy current-induced artifacts (ghosting and geometric distortions) in high gradient strength diffusion MRI (dMRI).

**Methods:** A dynamic field camera with 16 NMR field probes was used to characterize eddy current fields induced from diffusion gradients for different gradients strengths (up to 300 mT/m), diffusion directions, and shots in a 3D multi-shot EPI sequence on a 3T Connectom scanner. The efficacy of dynamic field monitoring-based image reconstruction was demonstrated on high-resolution whole brain ex vivo dMRI. A 3D multi-shot image reconstruction framework was informed with the actual nonlinear phase evolution measured with the dynamic field camera, thereby accounting for high-order eddy currents fields on top of the image encoding gradients in the image formation model.

**Results:** Eddy current fields from diffusion gradients at high gradient strength in a 3T Connectom scanner are highly nonlinear in space and time, inducing high-order spatial phase modulations between odd/even echoes and shots that are not static during the readout. Superior reduction of ghosting and geometric distortion was achieved with dynamic field monitoring compared to ghosting approaches such as navigator- and structured low-rank-based methods or MUSE, followed by image-based distortion correction with eddy. Improved dMRI analysis is demonstrated with diffusion tensor imaging and high-angular resolution diffusion imaging.

**Conclusion:** Strong eddy current artifacts characteristic of high gradient strength dMRI can be well corrected with dynamic field monitoring-based image reconstruction, unlike the two-step approach consisting of ghosting correction followed by geometric distortion reduction with eddy.

## 1. Introduction

High-performance gradients have become increasingly prevalent in human MRI scanners as gradient technology and hardware advance, largely spurred by developments initiated for the Human Connectome Project and other major neuroimaging initiatives.^1–3^ The use of ultra-strong MR gradients in diffusion MRI (dMRI) enables stronger diffusion weighting per unit time compared to scanners with conventional gradients, thereby shortening the time need for diffusion encodings, increasing the SNR, or enabling diffusion imaging at shorter diffusion times.^4^ The advantages of ultra-strong diffusion gradient measurements for microstructural imaging of the human brain have been demonstrated by multiple research groups.^3,5–13^ See ^4^ for an excellent review of applications.

Maximizing the benefits afforded by high-performance gradient technology requires overcoming the technical challenges associated with strong gradients, including more pronounced eddy currents, concomitant field effects, and peripheral nerve stimulation.^2,14^ Strong diffusion-sensitizing gradients induce eddy currents in the scanner bore and produce spatiotemporal variations in the magnetic field that depend on the strength and direction of the diffusion-encoding gradients. Concomitant field effects and peripheral nerve stimulation have been addressed iteratively in the design and calibration of gradient hardware.^15–19^ However, approaches to addressing eddy current-induced artifacts, such as geometric distortions and blurring, have largely relied on image post-processing tools due to ease of use and wide accessibility of the relevant software tools. Current state-of-the-art methods such as the ‘eddy’ tool from the FMRIB Software Library (FSL) correct for eddy current-induced distortions by utilizing the redundancy in directionally-sampled diffusion data along a sphere to solve for a model of eddy current-induced magnetic fields up to third order.^20,21^ However, eddy assumes that a single deformation field can explain and correct for all geometric distortions without accounting for the nonlinear temporal disturbance of the magnetic field induced by eddy currents during the image readout.^22–24^ Other potential drawbacks include failure in image registration convergence due to residual ghosting or insufficient SNR,^25^ and the introduction of blurring due to image resampling.

As the eddy current amplitude increases with the strength of the applied gradients, eddy currents induced by strong diffusion-sensitizing gradients can perturb the image encoding gradients.^2^ Previous work has shown that the lifetimes of eddy currents in high-gradient dMRI can be long-lasting and even exceed the standard echo planar imaging (EPI) readout duration.^26^ These effects are exacerbated for segmented acquisitions and can result in strong geometric distortions and ghosting artifacts that vary with the diffusion encoding direction.^23,27–32^ Existing methods that are effective for correcting ghosting artifacts observed with conventional gradient strengths^33,34^ underperform for multi-shot EPI dMRI experiments using strong diffusion-sensitizing gradients due to the complex nonlinear spatiotemporal magnetic field evolution incurred by the induced eddy currents.^35^ The residual ghosting artifacts bias subsequent dMRI analyses in a diffusion-direction dependent manner.^35^ Advanced k-space reconstruction methods that simultaneously handle nonlinear spatial phase variations between odd and even echoes and shots within an undersampled image reconstruction framework can effectively mitigate ghosting artifacts in high-gradient strength dMRI.^35,36^ Nevertheless, these methods do not explicitly incorporate information on the spatiotemporal phase evolution due to eddy currents in the image reconstruction model. They may underperform under certain circumstances, e.g., when the undersampled reconstruction problem becomes harder to solve due to increasing number of shots but limited coil sensitivity encoding in highly segmented, high-spatial resolution EPI acquisitions.

Magnetic field monitoring represents an alternative approach which directly measures and accounts for the nonlinear spatial and temporal evolution of eddy currents in the image reconstruction. Dynamic field monitoring has been validated for geometric distortion correction in single-shot EPI dMRI at 3T and 7T with b-values up to 2,000 s/mm^2^.^22,23^ In this work, we aim to demonstrate the efficacy of dynamic field monitoring^25,37^ for simultaneously correcting ghosting and geometric distortion artifacts in the challenging context of 3D multi-shot EPI, high-resolution dMRI using high gradient strengths and b-values up to 300 mT/m and 10,000 s/mm^2^, respectively. We demonstrate the benefits of dynamic field monitoring for ex-vivo whole human brain diffusion imaging, which requires strong diffusion-sensitizing gradients and segmented acquisitions^38^, e.g., 3D multi-shot EPI, to achieve sufficient diffusion contrast and SNR due to reduced diffusivity and shorter T2 relaxation times of fixed tissue. In this regime, eddy current artifacts can pose a significant barrier to achieving the high spatial resolution and image quality required to fully leverage the potential of this modality for mapping human brain structure at the mesoscopic scale.^38–42^

We used a dynamic field camera with 16 ^1^H NMR field probes distributed on the surface of a sphere to measure the actual phase evolution during the image readout of a 3D multi-shot EPI sequence on a 3T Connectom scanner. We provide evidence that eddy current fields induced by diffusion gradients with maximum strengths up to 300 mT/m are highly nonlinear in space (up to third order), persist for most of the readout period, and give rise to phase accrual terms that are also highly nonlinear in time, with different time course depending on the spatial order and diffusion direction. We adapt the multi-shot image formation model by incorporating the measured field evolution in each shot’s image encoding matrix. Solving this generalized image reconstruction problem allows us to reconstruct ghosting, distortion and blurring-free images. We show that this single-step approach outperforms two-step pipeline that consist of ghosting correction methods (navigator-based, structured low-rank-based approaches, or MUSE) followed by image-based geometric distortion correction using eddy. ^21^

## 2. Methods

### 2.1 Nonlinear phase modeling during image encoding

The phase evolution *ϕ*(*r,t*) of the transverse magnetization *M_xy_*(*r*) can be expressed at any point in space ***r*** = (*x, y, z*) and time *t* as

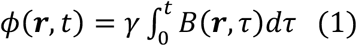

with *B*(*r, τ*) representing the magnitude of the main magnetic field and *γ* the gyromagnetic ratio.^43^ The k-space data, *y_p_* (*t*), received in the p-th coil (of a total of P coils), is related to *M_xy_* (*r*) (the image) by the following integral

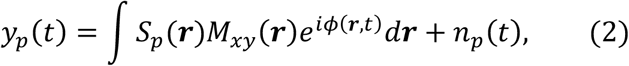

where the integration domain is the volume excited with the RF magnetic field (B1), *S_p_* (*r*) is the coil sensitivity profile of the p-th coil, and *n_p_* (*t*) denotes the noise in the measurements.^43^ In MR image encoding, the spins are spatially encoded with a time-varying linear magnetic gradient field oriented along the B° field. The phase *ϕ*(***r**, t*) is thus linear in space and *M_xy_* (*r*) in Eq.(2) can be reconstructed with a Fourier transform.^43^ In practice, nonlinear fields perturbations interfere with the image encoding gradient, rendering the linear phase assumption invalid.^44^ If Fourier reconstruction is used, artifacts will appear, most notably geometric distortions and ghosting in diffusion-weighted EPI acquisitions.^25,30,33,45^ Provided the mathematical expression of *ϕ*(*r,t*) is known, including any nonlinear phase perturbations, accurate reconstruction of the image *M_xy_* (*r*)is still possible if the reconstruction problem (Eq. (2)) is treated as a general inverse problem.^25,46,47^ This section describes the nonlinear model for *ϕ*(***r**, t*) that will be later used in the 3D multi-shot reconstruction framework applied to high gradient strength dMRI ex vivo brain data.

Nonlinear field perturbations can be classified as static or dynamic.^37^ An example of static magnetic field perturbations include those generated by magnetic susceptibility effects from the sample, resulting in off-resonance effects ΔB_0_(***r***).^45^ Dynamic magnetic field perturbations include those generated by changes in the gradient magnetic fields and will depend on the pulse sequence. Concomitant fields (also known as Maxwell fields) and eddy current fields are the most important contributions to dynamic nonlinear field changes.^25^ Concomitant fields can be derived from the Maxwell equations and predicted analytically from the applied gradient magnetic field.^48–51^ Characterizing the eddy current fields is arguably more challenging. Eddy currents generated in the main magnet and gradient system can be characterized using network analysis.^52–54^ The spatial distribution, i.e., the functional form of the magnetic field produced by eddy currents may be approximated using a linear combination of spatial functions with known closed-form expressions.^55,56^ The representation’s accuracy is contingent on the mathematical assumptions that can be made about the eddy current fields. A common assumption is that the induced eddy current fields are smooth in space and thus can be approximated by smooth basis functions.^25,37^ Based on this assumption, several investigators have suggested using regular solid spherical harmonics as the basis functions to express the eddy current fields.^25,37,55,56^ Following previous work, we assume that a solid spherical harmonics expansion up to a third order is sufficient to describe the eddy current fields in the current framework.^22,25^ The spherical harmonic expansion includes time-varying coefficients that reflect the temporal variation of the eddy current fields. Because the integral in Eq.(1) is a linear operator with respect to *B*(*r,τ*), the integral and summation of the integrand can be interchanged, and the phase distribution of eddy current fields can also be written as a linear combination of solid spherical harmonics. Therefore, we adopt the following spatiotemporal phase model during the image readout of the 3D diffusion-weighted multi-shot EPI used in this study:

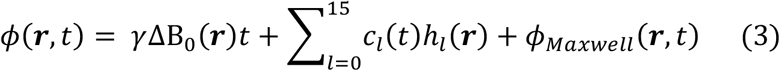

where *h_l_*, (*r*) are the real-valued solid harmonics up to third order (see Table S1 in the supplementary file), *c_l_*, (*t*) the corresponding time-varying expansion coefficients, and *ϕ_Maxwe_*(***r**, t*) is the phase accrual due to concomitant fields. The k-space trajectories, *k_x_*(*t*), *k_y_*(*t*), and *k_z_*(*t*), which are driven by the image encoding gradients, are accounted for in the coefficients *c_l_*,(*t*) (*l* = 1,2,3) corresponding to the solid harmonics *h*_1_***r***) = *x*, *h_2_*(***r***) = *y*, and *h*_3_(***r***) = *z*, respectively. Note that the temporal coefficients *c_l_*, (*t*) (*l* = 1,2,3) also describe the linear components of eddy current fields. We must estimate ΔB_0_(***r***) and *c_l_*, (*t*) to properly determine the nonlinear phase model of Eq.(3). ΔB_0_(***r***) is calculated with a multi-gradient echo-based acquisition in this study.^57^ We measure *c_l_*, (*t*) via dynamic field monitoring as described next.

### 2.2 Dynamic field monitoring: Acquisition set-up, 3D diffusion-weighted multi-shot EPI sequence, and signal processing

The magnetic field was monitored during a 3D diffusion-weighted multi-shot EPI sequence with Stejskal-Tanner encoding^40^ (Figure 1A) using a dynamic field camera^58^ (Skope MR Technologies Inc., Zurich, Switzerland) in a 3T Connectom Scanner (Siemens, Erlangen, Germany). The dynamic field camera measures the free induction decay (FID) signals (at a sampling rate of 1MHz) from 16 ^1^H-based field probes distributed evenly on the surface of a 18 cm diameter sphere.^58,59^ The NMR probes are smaller than the spatial resolution of the EPI acquisition and can thus be considered as point samples.^59^

**Figure 1.**
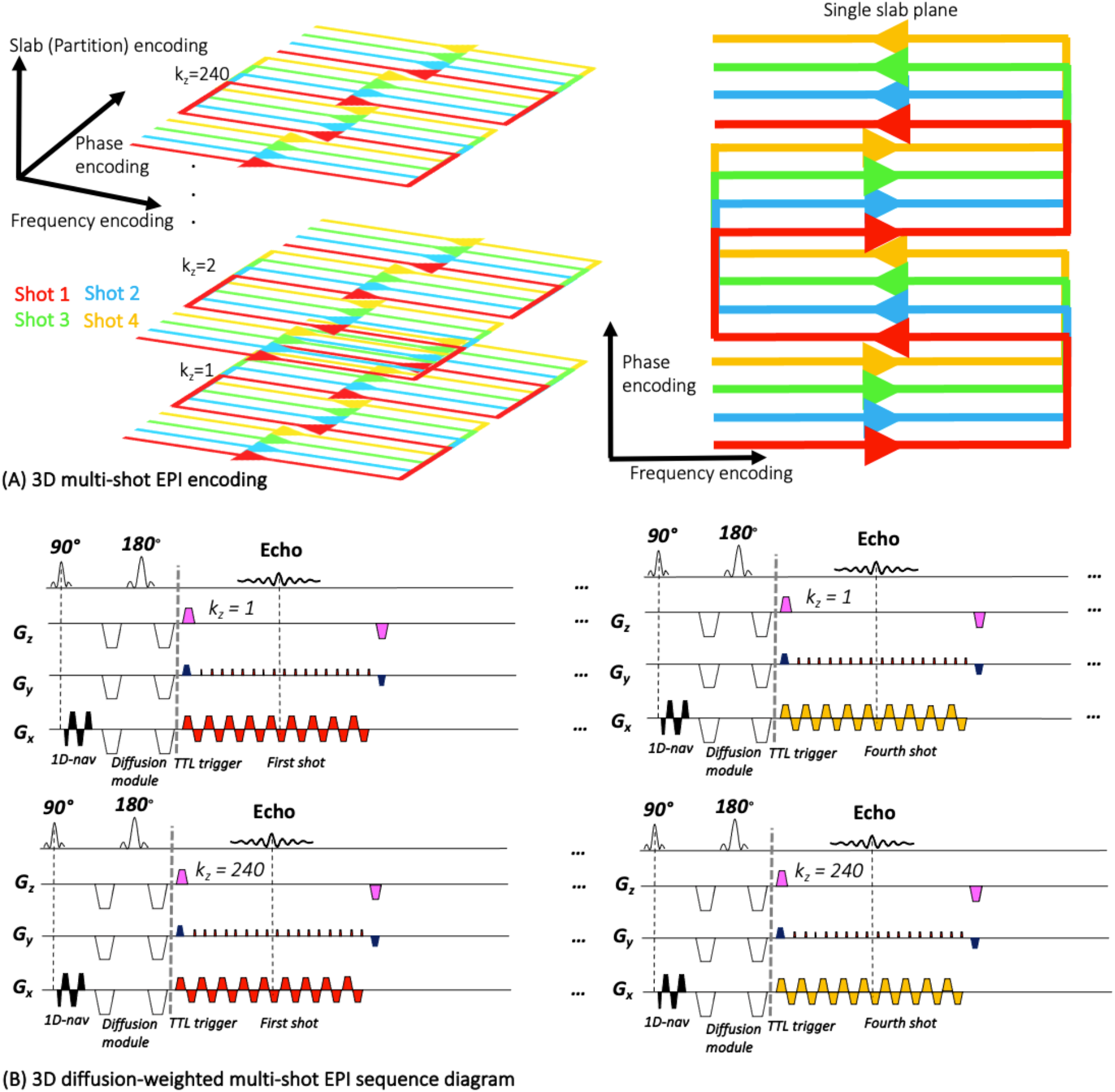
Representative k-space and sequence diagrams for the 3D diffusion-weighted multi-shot EPI sequence used in this work. (A) K-space diagram outlining 3D encoding along the frequency, phase, and slab (partition) directions. Separate shots are color-coded in red, yellow, green, and blue for this representative four-shot experiment. (B) 3D diffusion-weighted multi-shot EPI sequence diagram with Stejskal-Tanner diffusion encoding. TTL triggers (denoted in pink) were placed immediately after the diffusion module to signal the start of field monitoring.

The phase accrual due to Maxwell terms must be calculated before determining *c_l_*, (*t*). A first-order solid spherical harmonic model was fitted to the phase of the signals obtained from the 16 NMR field probes, from which the image encoding gradients were measured and then used to derive the concomitant fields terms based on equations for symmetric coils, as in the case of the Connectom whole-body gradient coil.^15^ After performing the integral in Eq.(1), the term *ϕ_Maxwe_*, (***r**, t*) was then subtracted from the 16 acquired phase signal. The expansion coefficients *c_l_*, (*t*) were then determined by fitting the phase signals to a linear combination of third-order solid spherical harmonics using a linear least-squares (LLS) estimator.^37^

A Transistor-Transistor-Logic (TTL) signal was used as a trigger to start the dynamic field measurements immediately after the diffusion module, precisely 400 microseconds after the end of the diffusion gradient waveform. The TTL trigger was repeated every TR (for each shot and partition). See Figure 1B for a schematic diagram of the 3D diffusion-weighted multi-shot EPI sequence including the trigger. A common time base was established between the field camera and MRI system, accounting for receiver chain delays and different sampling rates. The two systems were synchronized by locking the camera’s reference oscillator onto the MRI system’s reference clock. An amplitude-modulated test signal was transmitted into both receive chains during a synchronization scan to calibrate the delay. The sampling rate difference was rectified by resampling the observed k-space trajectory to match the bandwidth of the raw MR imaging data, which is typically acquired at lower bandwidth.

### 2.3 Dynamic field monitoring-based multi-shot image reconstruction

Let *ϕ_k_z_s_* (***r**,t*) represent the phase evolution measured with the field camera during the image readout of the kz partition or slab encoding index (from a total number of Kz partitions or slabs) and shot index.s’ (from a total number of shots *S*). Let *y_k_z_,s,p_*(*t*) denote the corresponding k-space data acquired with the p-th coil. To develop the multi-shot image reconstruction framework, we discretized the continuous forward model of Eq. (2) in space and time. We used a spatial grid 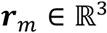 consisting of *M* points that covered the entire field of view (FOV) with grid spacing set to the nominal resolution. The temporal grid *t_n_* contains *N* time points for each read-out, with time intervals equivalent to the analog to digital converter (ADC) dwell time. We call ***y**_k_z_,s,p_* = [*y_k_z_,s,p_*(*t*_1_),…, *y_k_z_,s,p_*(*t_N_*)]^*T*^ the k-space data vector containing time samples of *y_k_z_,s,p_*(*t*) and ***E**_k_z_,s_* the image encoding matrix associated with *k_z_* partition and shot index *s*, whose elements are defined as {***E**_k_z_,s_*} = *e*^*iπ_;k_z_,s_*(***r***_m_,*t_n_*^) (*N* < *M*).

The forward model that relates the image ***x*** = [*M_xy_* (***r**_1_*),…, *M_xy_* (***r**_M_*)]^*T*^ with the measurements *y_k_z_,s,p_* is then

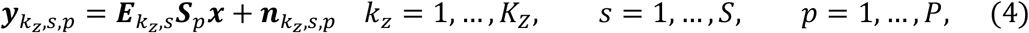

with ***S**_p_* a diagonal matrix with elements {***S**_p_* (***r**_m_*)}, and *n_k_z_s,p_* representing complex Gaussian noise. We seek to recover the image ***x*** from the measured signal ***y**_z,s,p_*. For the fully-sampled 3D multi-shot acquisitions considered in this study, this inverse problem is well-posed. While the matrices ***E**_k_z_,s_* are rank-deficient, the k-space trajectories of each shot and partition are complementary, such that the entire k-space is covered, satisfying the Nyquist criterion. For fully-sampled reconstruction problems, and provided ***S**_p_* are known, a conventional least squares estimator with Tikhonov regularization is known to yield satisfactory image reconstruction:^25,43,46,60^

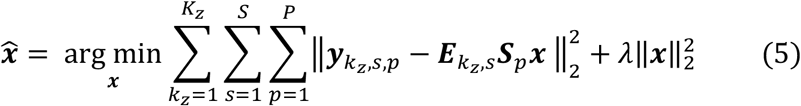

where *λ* representing the Tikhonov regularization parameter. Coil sensitivities ***S**_p_* were determined using the ESPIRiT technique from a GRE acquisition taken prior to the dMRI acquisition.^61^ The phase difference of two GRE images was utilized to estimate ΔB_0_(***r***).

The solution 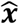 can be obtained in closed form with the Moore-Penrose inverse matrix, which models the linear relationship between ***x*** and measurements ***y**_k_z_,s,p_*.^35^ However, this strategy was not feasible to achieve in practice because of the large dimensionality of the data at hand. The following assumption made the problem tractable. As the identical diffusion gradient waveform (for a chosen diffusion direction) is played out every slab encoding, *k_z_*, we assumed that most of the differences of *ϕ_k_z_,s_*(***r***, *t* along *k_z_* were attributed to *c*_3_(*t*), associated to *h*_3_(***r***) = *z* (nominal k-space trajectory along *z* and linear eddy current fields). The temporal differences in the remaining spherical harmonic coefficients across partitions were assumed to be insignificant compared to intra-partition temporal variations, e.g., phase fluctuations during the image readout and between shots, which are the most prominent source of ghosting and geometric distortions. We have used this assumption before with satisfactory results.^35^ We then applied a non-uniform inverse Fourier Transform along the partition direction *k_z_*,^62^ and treated the problem as a 2D multi-shot reconstruction approach for each slice index *z*. We adjusted the image encoding matrix ***E**_k_z_,s_* by: 1) removing the contribution of *c*_3_(*t*) to the phase model, and 2) modifying the grid *r_m_* to account for a fixed slice *z*. We then solved a similar Tikhonov regularization problem similar to Eq.(5) for each *z* partition. In the Discussion section, we explain the implications of this assumption and suggest future extensions that can handle the exact 3D reconstruction problem efficiently.

The LSQR algorithm was used to find the solution of Eq.(5) iteratively.^63^ Convergence was achieved after 10 iterations and took approximately 20 min per slice. The tolerance between iterates was set to 10^-10^. The code was written in MATLAB 2019b. The Tikhonov regularization parameter was chosen empirically as done in similar work. ^64^ We found *λ* = 10^-6^ to be a reasonable choice, providing satisfactory results for all three post-mortem brain specimens imaged in this study. It should be noted that the choice of *λ* does not affect the extent of ghosting and geometric distortion correction and only affects the SNR of the reconstructed images. Other approaches to selecting this parameter and potential strategies to compensate for reduced SNR are described in the Discussion section.

### 2.4 Experiments

#### 2.4.1 Experiment 1: Characterization of eddy current fields induced by diffusion-sensitizing gradients

We first studied the spatiotemporal behavior of eddy current fields induced by the diffusion-sensitizing gradients on the 3T Siemens Connectom scanner for different diffusion directions and gradient strengths. The phase evolution was measured during the 3D diffusion-weighted multishot EPI sequence shown in Figure 1B with 1 b0-image and 6 non-collinear diffusion directions (see legend of Figure 2) for b = 1,500 s/mm^2^ (G_max_ = 94 mT/m), b =4,000 s/mm^2^ (G_max_ = 171 mT/m), b = 6,000 s/mm^2^ (G_max_ = 232 mT/m), and b = 10,000 s/mm^2^ (G_max_ = 277 mT/m), during an image readout length of 50 ms. Other sequence parameters were: TE/TR = 81/500 ms, FOV = 170 × 125 × 176 mm^3^, matrix size = 170 × 125 × 176, no acceleration, no partial Fourier (PF), EPI factor = 42, #Shots = 3, echo spacing (ESP) = 1.28 ms.

**Figure 2.**
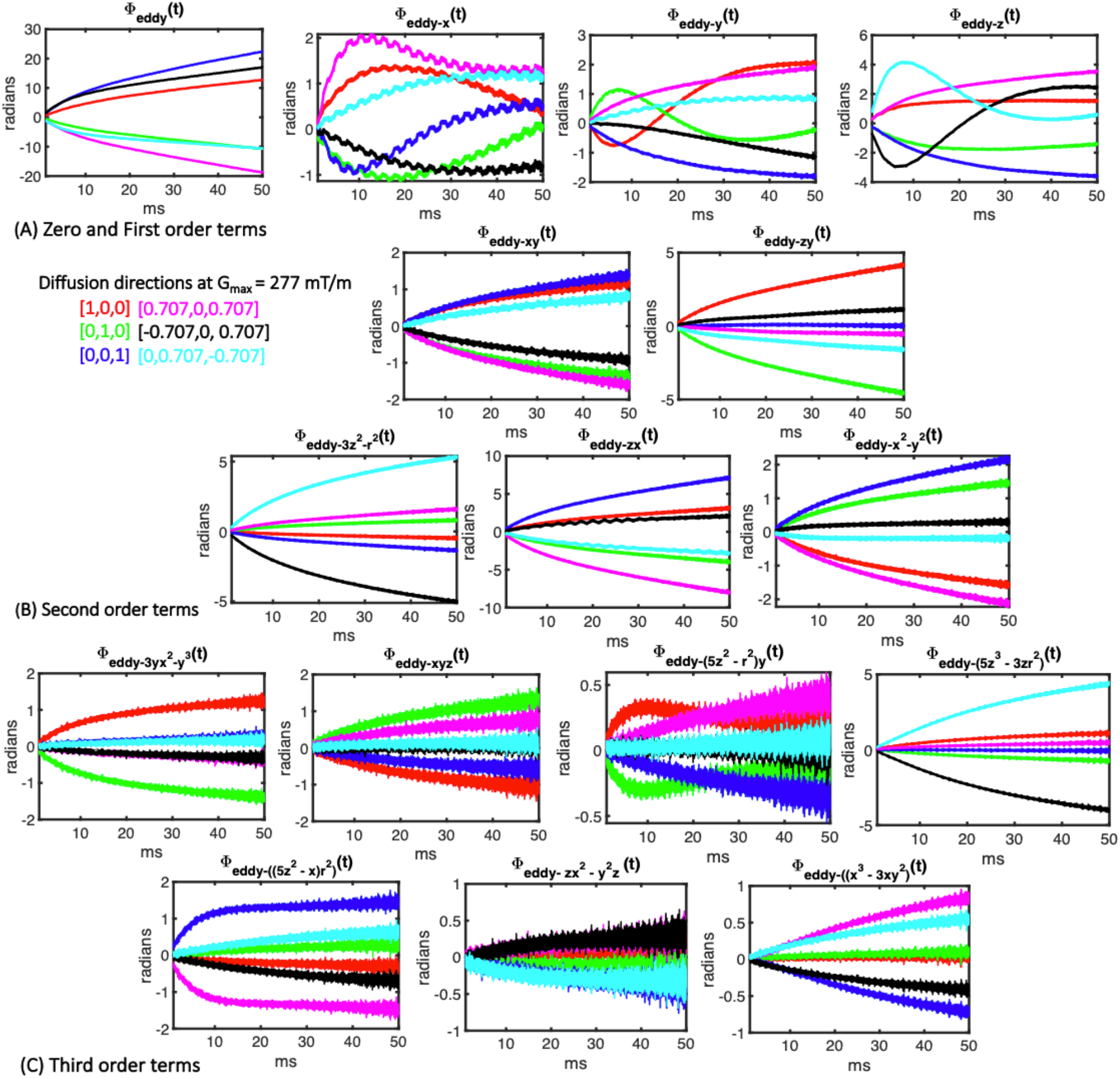
Spatiotemporal phase of the eddy current fields induced by diffusion gradients at G_max_ = 277 mT/m, for the first shot and kz close to the zero frequency, as expressed in a basis of solid spherical harmonics up to third order. The maximum phase accrual achieved within the FOV is shown in radians *ϕ_eady-h_l__*(*t*) for every harmonic *h_l_*,(***r***).

We subtracted the spherical harmonics coefficients of the phase evolution during the image encoding of the b0-image from those during the image encoding of the diffusion-weighted images. If heating effects are ignored, the difference coefficients, *k_eddy-l_*, (*t*), capture only the phase variations reflecting the eddy currents induced by the diffusion-sensitizing gradients.^11,47^ For each gradient strength, diffusion direction, shot and partition *k_z_*, we calculated *ϕ_eddy-h_l__*(***r**,t*) = *k_eddy-l_* (*t*)*h_l_*, (***r***), the eddy current induced phase evolution expressed by the l-th spherical harmonic *h_l_*, (***r***). We also analyzed the variation of eddy current fields among shots and for odd and even EPI echoes during the readout.

#### 2.4.2 Experiment 2: Correction of ghosting and geometric distortions induced by eddy currents in high gradient strength dMRI experiments on ex-vivo human brain samples

We next investigated the ability of dynamic field monitoring to simultaneously correct for ghosting and geometric distortion artifacts induced by eddy currents from strong diffusion-sensitizing gradient strength in dMRI scans of three post-mortem human brains. All samples were scanned in the 3T Siemens Connectom scanner using a 3D diffusion-weighted multi-shot EPI sequence (Figure 1(B)). A dedicated 48-channel receive array coil constructed specifically for high-sensitivity mesoscopic dMRI of whole human brain specimens was used for MR signal reception.^65^ The magnetic field was monitored immediately after each ex-vivo brain scan using the same sequence and protocol parameters. Field monitoring was performed using the dynamic field camera (Skope, Zurich, Switzerland), which was placed at the gradient coil isocenter. All brains were positioned in the dedicated whole-brain ex vivo coil, as shown in Figure S1 of the supplementary file. Scans were conducted with the approval of MGB Institutional Review Board. A description of the three post-mortem brain specimens and acquisitions is provided below and in Table 1.

1. <u>Ex vivo brain 1:</u> Whole fixed human brain from a male who died of non-neurological causes. The excised brain was placed in fixative (10% formaldehyde) for 90 days and transferred to a paraformaldehyde-lysine-periodate solution for long-term storage.
2. <u>Ex vivo brain 2:</u> Whole fixed human brain from a female who died from traumatic brain injury. The specimen was fixed in formaldehyde and packed in Fomblin (Ausimont USA Inc.) for the scan. A middle coronal slab was scanned using a small field of view.
3. <u>Ex vivo brain 3:</u> Left hemisphere of human brain from a male who died of non-neurological causes. The excised brain was placed in fixative (10% formaldehyde) for longer than 90 days and transferred to a paraformaldehyde-lysine-periodate solution for long-term storage.

**Table 1.**
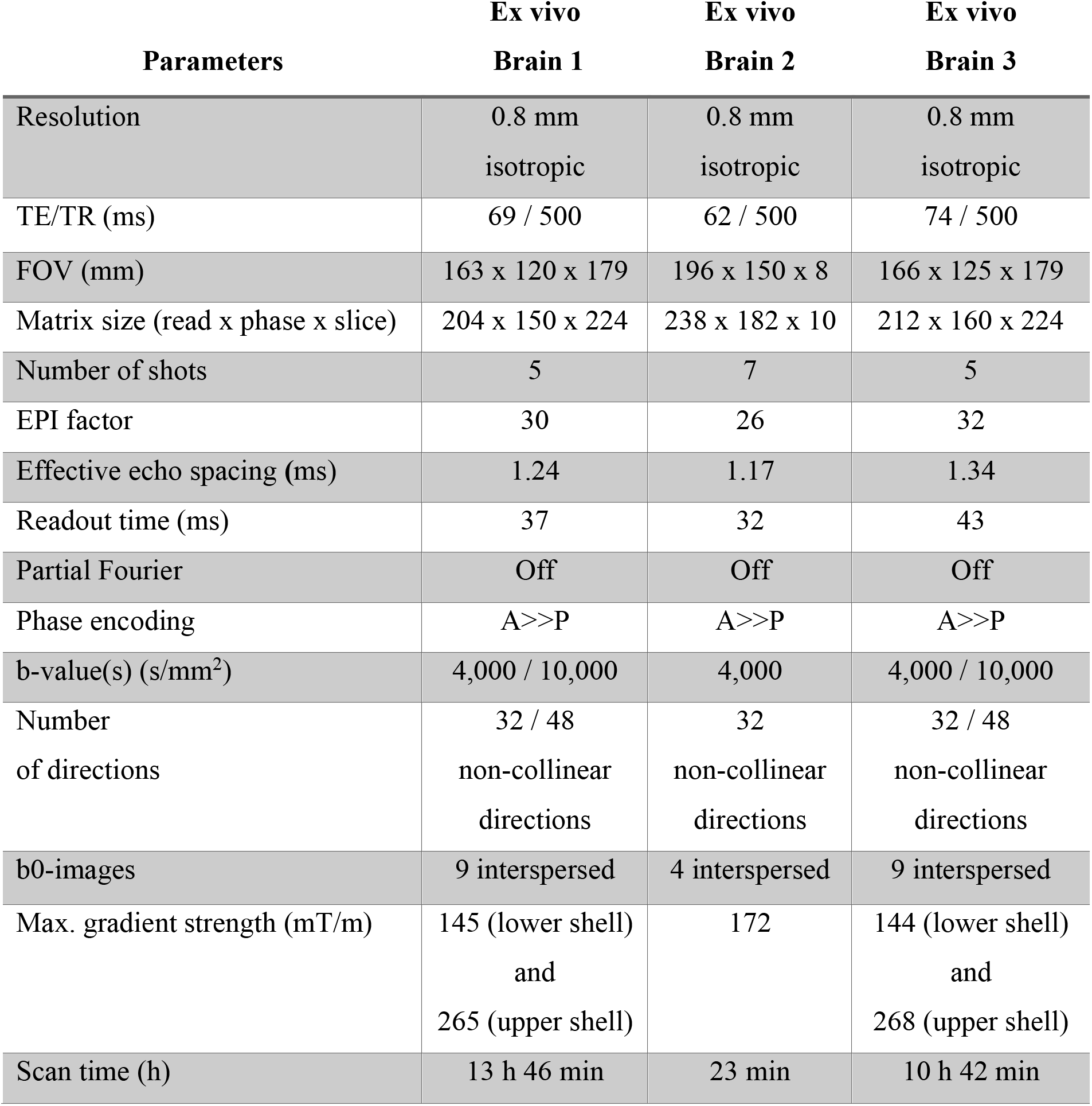
Acquisition parameters for the experiments on the three ex vivo human brains.

The images were reconstructed using the magnetic field trajectories obtained from dynamic field monitoring. For comparison purposes, image were also reconstructed using other ghosting correction techniques, followed by geometric distortion correction with the FSL’s tool ‘eddy’. The ghosting correction methods implemented were:

1. Conventional one-dimensional linear phase correction method (1D LPC) to account for odd/even phase modulations at each shot.^33^ Corrected k-space lines of each shot were combined to get a fully-sampled k-space dataset. An image was then reconstructed with 3D SENSE.^66^ The one-dimensional navigator (three k-space lines) was played out for every shot, diffusion direction and k_z_ partition.
2. The multi-shot technique 2D MUSE ^34^, which account for nonlinear phase variations among shots. The phase modulation between odd and even lines was accounted for with one-dimensional linear phase correction. 2D MUSE was applied right after a non-uniform inverse Fourier transform along the slab encoding.
3. Our previously published method that extends the structured low-rank matrix-based method^36^ to a multi-shot acquisitions and handles both nonlinear spatial variations in phase between odd and even echoes and between shots simultaneously (SLM-based method).^35^ As with 2D MUSE, the k-space data was first non-uniform inverse Fourier transformed along the slab encoding. Shots and odd/even images were combined with rSoS.

The reduction of ghosting artifacts was assessed visually and quantitatively with the ghost-to-signal ratio (GSR) metric. This metric quantifies the level of ghost-only signal with respect to the signal in the rest of the sample where ghosting has been reduced to below a minimum threshold. Although the lack of ground truth is a known issue in the EPI ghost-correction literature, the GSR metric can provide insight into the relative degree of ghosting reduction without the need for a ghost-free reference acquisition, which may be challenging to achieve for a given set of acquisition parameters. As such, the GSR has been used extensively in other work to quantify the degree of ghosting reduction. ^35,67–69^ For the ex vivo human brain experiments, the mean signal was calculated over all gray and white matter voxels. Ghost-only signal was calculated in the background voxels with non-zero coil sensitivity. The same coil sensitivities that were employed to reconstruct images with dynamic field monitoring ***S**_p_* were used for the implementation of the three ghosting correction methods.

FSL’s ‘eddy’ algorithm was run on the image reconstructed with the ghosting reduction methods described previously. Spline interpolation was used for both registration and estimation steps. A cubic model was chosen for the eddy current deformation field. To ensure convergence, the number of iterations was set to 30 (compared to 5 by default). The FSL ‘topup’ function was to correct for susceptibility-induced image distortion using an additional b0-image acquired with reverse phase encoding. The quality of geometric distortion correction was assessed both visually and quantitatively. We quantified the image sharpness after eddy current correction using image-based registration (‘eddy’) and dynamic field monitoring-based image reconstruction. We used two non-reference image blur metrics: the cumulative probability of blur detection (CPBD) metric^70^ and the perceptual image sharpness (PSI) metric based on local edge gradient analysis.^71^ Both image blur metrics are defined in the interval (0,1). The sharper the images look, the higher the CPBD and PSI values are.

To investigate the downstream effects of ghosting and geometric distortions on the estimation of dMRI metrics, we performed diffusion tensor imaging (DTI) with the linear least squares (LLS) estimator, as implemented in the FSL tool ‘*dtifit*’. In addition, fiber ODFs were estimated with the multi-shell, multi-tissue constrained spherical deconvolution method implemented in the MRtrix3 software toolbox using the default parameters.^72,73^ After visual assessment, we calculated fractional anisotropy (FA) and mean diffusivity (MD) values estimated in selected white matter regions: corpus callosum, left corona radiata, and left temporal lobe. As in similar studies,^35,40^ region-of-interest (ROI) masks were hand-drawn for the three ex vivo brains.

## 3. Results

### 3.1 Experiment 1: Characterization of eddy current fields induced by diffusion-sensitizing gradients

Figure 2 shows the temporal evolution of the phase of the eddy current fields induced by diffusion-sensitizing gradients of G_max_ = 277 mT/m for a fixed partition index *k_z_* and the first shot of a three-shot EPI acquisition. For each spherical harmonic, *h_l_*,(***r***), we report the maximum phase accrual (in radians) achieved within the whole FOV, *ϕ_eddy-h_l__* (*t*). As observed, substantial phase contributions do exist from higher-order eddy current fields along *x*, *y*, and *z*. Note that the phase accrual from third-order spherical harmonics (Figure 2C) were, for some diffusion directions, comparable to first-order (Figure 2A) eddy current fields, see *ϕ*_*eddy*-*5z^3^*-*3zr^2^*_(*t*) or *ϕ*_*eddy*-(*5z^2^*-*x*)*r*^2^)_ (*t*). The observed temporal evolution of the phase induced by eddy current fields was highly nonlinear and differs across diffusion directions. The strongest phase variation was found in the linear components. Interestingly, *ϕ_eddy-h_l__*(*t*) did not reach a constant value in most cases, indicating that some eddy current fields had lifetimes on the order of milliseconds and could interfere with the image encoding gradients, resulting in ghosting and geometric distortion artifacts, as demonstrated in Experiment 2. Similar findings were obtained for other gradient strength settings, as shown in Figures S2 and S3 in the supplementary file. As expected, the higher the gradient strength, the greater the phase accrual (in radians), with the temporal profile of *ϕ_eddy-h_l__*(*t*) remaining unchanged.

Eddy current fields from diffusion-sensitizing gradients create phase modulations between odd and even EPI echoes are nonlinear in space and time and contribute to Nyquist ghosting. To demonstrate this, we calculated for each shot, diffusion direction, and partition k_z_, the increment of phase between k-space lines (u), Δϕ*_eddy-hl_*, (***r**, u*). Specifically, we computed the phase difference between adjacent k-space lines as measured in the middle of the k-space lines (every ESP = 1.28 ms). The time-varying coefficients of the spherical harmonic expansion of that difference Δϕ*_eddy-hl_*, (***r**, u*), Δk_*eddy-hl*_,[*u*], are displayed in Figure 3 (only components up to second order are shown for brevity). Note that the phase difference between adjacent k-space lines for the first-order spherical harmonics terms (Δk*_eddy-x_* [*u*], Δk_*eddy-y*_ [*u*], and Δk_*eddy-z*_ [*u*], with units in rad/m) can be interpreted as unwanted shifts of the nominal k-space trajectory due to eddy currents, with Δk_*eddy*_ [*u*] representing the phase offset, when moving from k-space line *u*-1 to *u*.As observed, eddy current fields introduce phase offset, k-space shifts, and spatially nonlinear phase increments between k-space lines that vary with the diffusion-encoding direction, potentially causing ghosting artifacts that differ across diffusion-weighted images. In general, the magnitude of the k-space shift appears greater at the beginning of the read-out and seems to stabilize during the last part of the read-out, although for certain terms, e.g., Δk_*eddy-xy*_ [*u*], the amplitude of this difference term appears to increase during the last portion of the readout. An important implication of this finding is that ghosting correction methods that use navigators which do not traverse the entire set of k-space lines may not be completely capturing eddy current-induced phase modulation, resulting in residual Nyquist ghosting.

**Figure 3.**
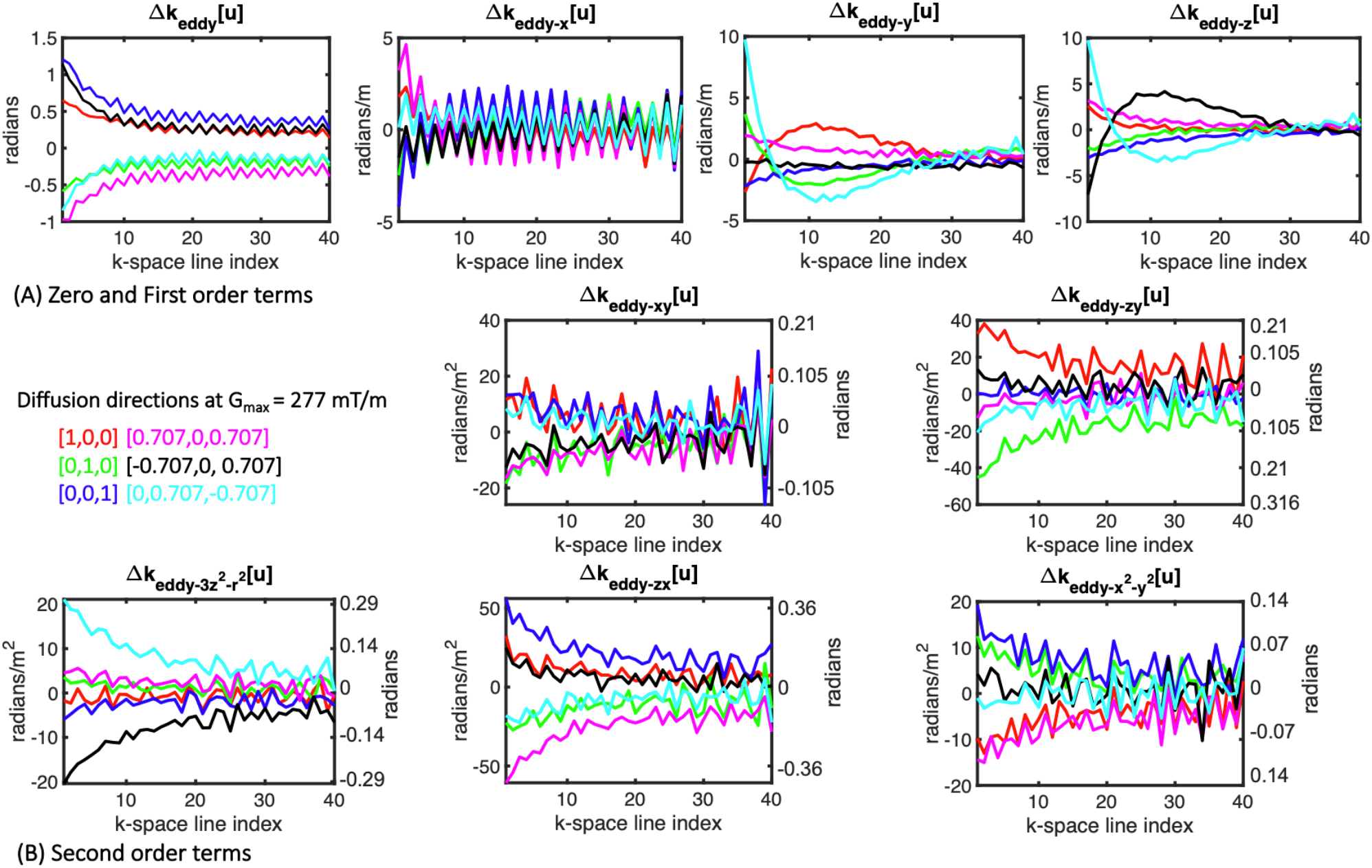
Phase modulation between odd and even EPI lines introduced by eddy current fields induced by diffusion gradients with G_max_ = 277 mT/m (first shot and identical k_z_ as in Figure 2). For brevity, only zero (radians), first-order (rad/m) (A), and second-order terms (rad/m^2^) (B) are shown. Maximum phase accrual within the FOV is also reported for the second-order terms (right axis). Note that the phase modulation between consecutive k-space lines differs between diffusion-encoding gradient directions and is not constant during the readout period.

We also report the phase variation of eddy current-induced fields among EPI shots (see Figure S4). Though the amount of eddy current fields variation between shots is generally smaller than that between k-space lines, especially for second-order terms, non-negligible variation does exist and this should be accounted for in the implementation of ghosting correction methods. In some cases, e.g., see Δk_*eddy-z*_ [*u*] for direction (0, 0.707, −0.707), the phase of the diffusion-gradient induced eddy current fields vary more between shots than between odd/even EPI echoes.

### 3.2 Experiment 2: Correction of ghosting and geometric distortions induced by eddy currents in high gradient strength dMRI experiments on ex-vivo human brain samples

Figure 4 shows two coronal slices of the same diffusion-weighted volume from the ex vivo brain 1 dataset reconstructed with dynamic field monitoring-based image reconstruction, 1D LPC followed by 3D SENSE, 2D MUSE, and the SLM-based ghosting correction method.

**Figure 4.**
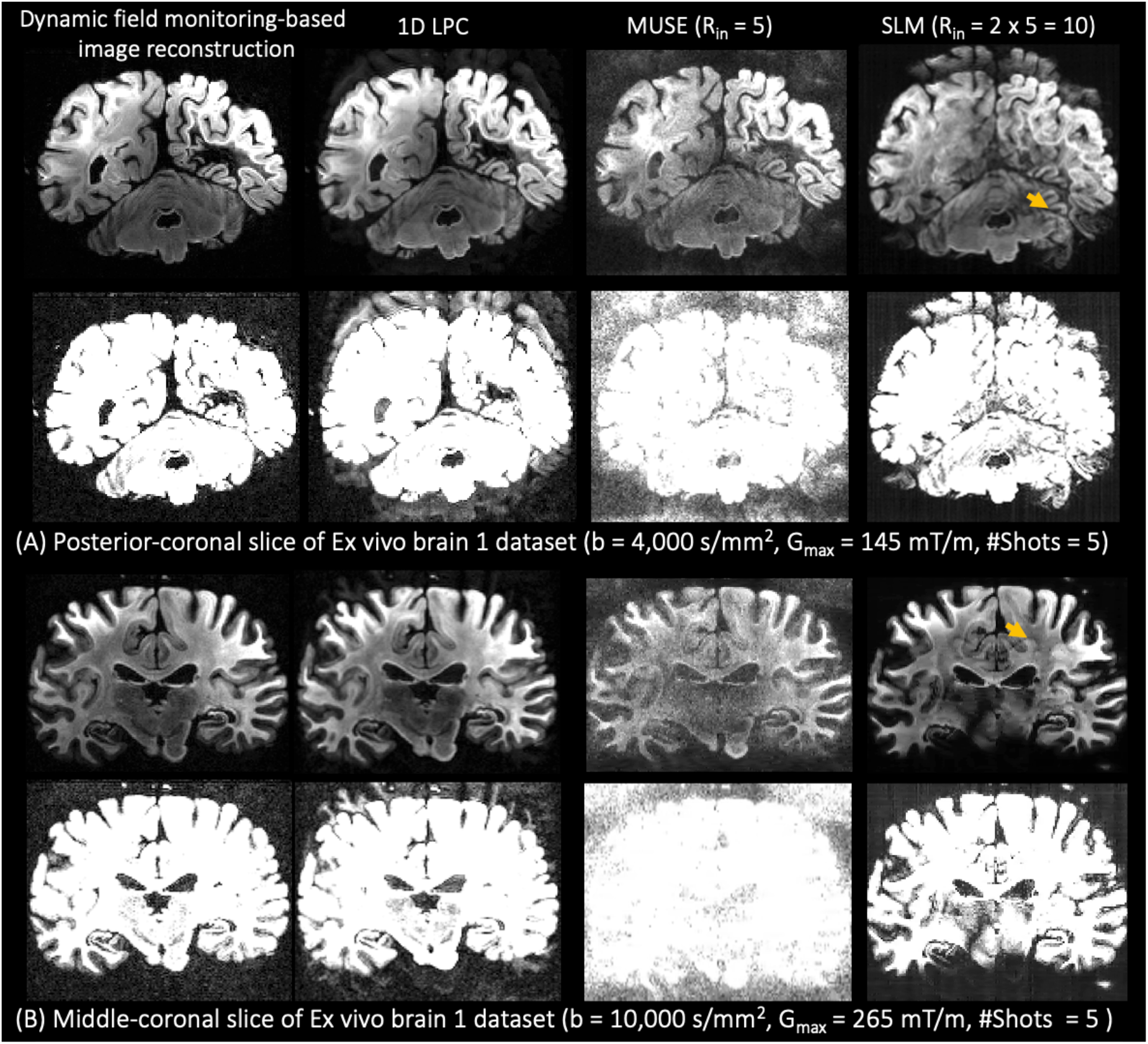
Two representative coronal slices from the ex vivo brain 1 dataset reconstructed with dynamic field monitoring-based image reconstruction, 1D LPC, MUSE, and SLM-based ghosting method at (A) G_max_ = 150 mT/m and (B) G_max_ = 270 mT/m. Both MUSE and the SLM-based method suffer from reconstruction artifacts (yellow arrow). Ghosting artifacts are reduced to a greater extent with dynamic field monitoring compared to 1D LPC.

Images are also presented with the intensity range limited to 20% of the maximum original value for better visualization of the ghosting artifacts. Results from the two other ex vivo datasets are provided in the supplementary file (Figures S5 and S6). Taken together, the images reconstructed across all three brains using MUSE and SLM-based approach suffer from reconstruction artifacts (see yellow arrow in Figure 4 for the SLM approach). The performance of the SLM-based method degrades when the number of shots increases, as the undersampled problem becomes harder to solve. Indeed, this method attempts to reconstruct, for each shot, a single image for each of the two k-space datasets that contain odd and even k-space lines only. Therefore, each k-space dataset is actually undersampled by a factor of two times the number of shots, i.e. *R*_in_ = 2 × 5 = 10, in this dataset. Our previous work demonstrated that the SLM-based approach can provide excellent ghosting reduction, handling nonlinear ghosting variation between shots and polarities in a four-shot 3D segmented EPI acquisition at similar spatial resolution (0.8 mm).^35^ But highly segmented acquisitions, as shown here, necessitate alternative solutions. Indeed, neither MUSE nor the SLM-based method could reconstruct acceptable images for the seven-shot EPI acquisition in the ex vivo brain 2 dataset (Figure S5).

Dynamic field monitoring and 1D LPC were able to reconstruct images without evident reconstruction artifacts in all three datasets. However, substantial ghosting was seen with 1D LPC. Note the remaining ghosting artifacts in the cortex or at the bottom of the cerebellum in Figure 4. Dynamic field monitoring achieves a remarkable reduction of ghosting artifacts, consistent with quantitative results. Figure 5 shows the GSR plotted as a function of the diffusion direction index. Compared to 1D LPC and 2D MUSE, dynamic-field monitoring-based image reconstruction yields a more than fivefold GSR improvement on average (MUSE’s GSR values are only presented for cases where MUSE was able to reconstruct an image, Figures 5A and 5B). Figures S3 and S4 showed that when G_max_ increases, so does the magnitude of the eddy current fields. As a result, the GSR for all methods increases when the gradient strength increases. In the case of dynamic field monitoring, that increase is only about less than 2%, as opposed to 8% and 15% for 1D LPC and MUSE.

**Figure 5.**
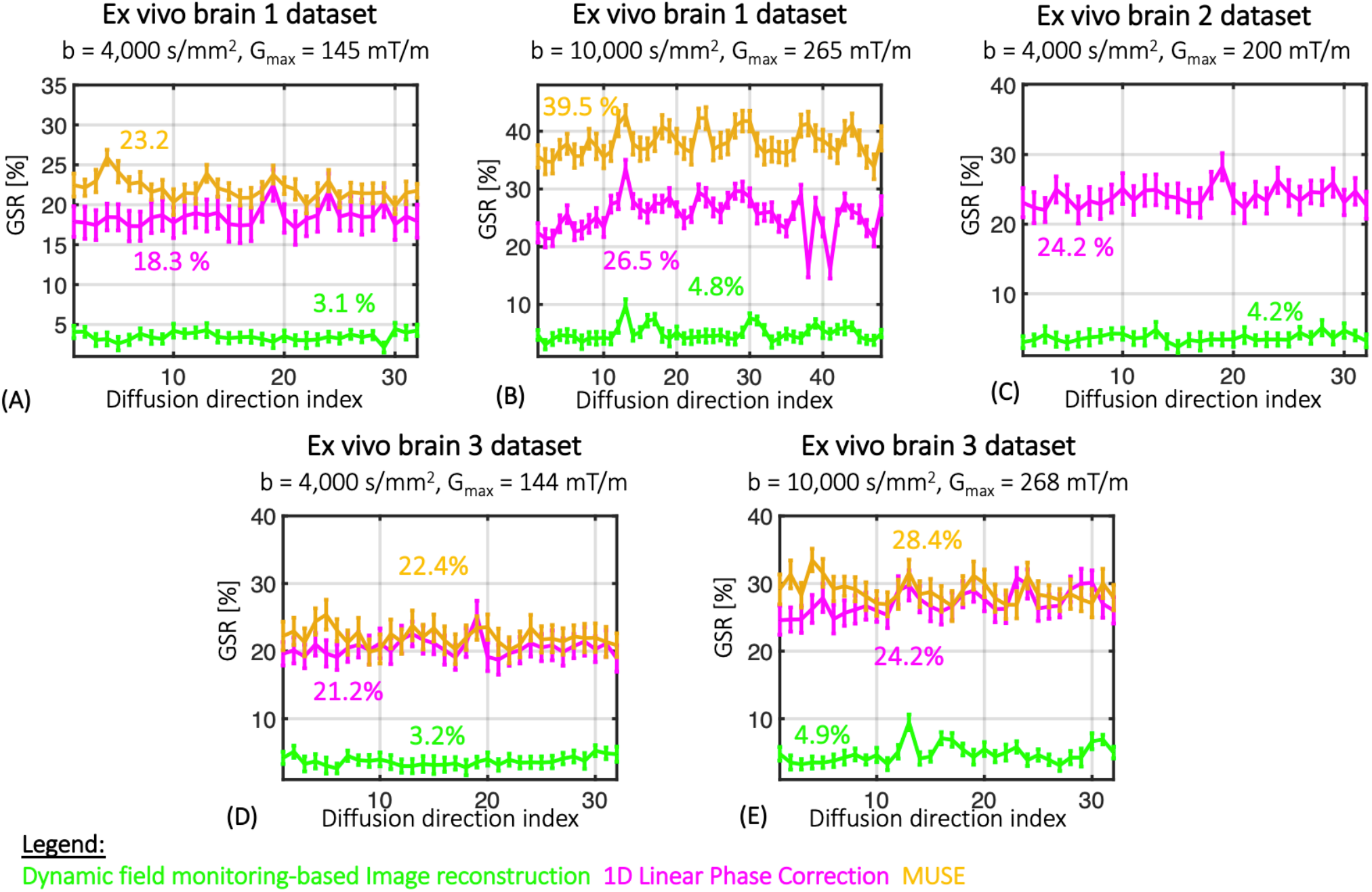
Ghosting-to-Signal Ratio (GSR) plotted as a function of the index of diffusion direction for dynamic field monitoring, 1D LPC, and MUSE reconstruction with (A) ex vivo brain 1 dataset at b=4,000 s/mm^2^ and G_max_ = 150 mT/m, (B) b = 10,000 s/mm^2^ and G_max_ = 270 mT/m, (C) ex-vivo brain 2 dataset at b = 4,000 s/mm^2^ and G_max_ = 200 mT/m, and ex vivo brain 3 dataset at (D) b=4,000 s/mm^2^ and G_max_ = 144 mT/m and (C) b =10,000 s/mm^2^ and G_max_ =268 mT/m. Note that MUSE was unable to reconstruct acceptable images for ex vivo brain 2 dataset and hence GSR is not shown.

Videos included in the supplementary file reveal that dynamic field monitoring outperforms image-based approaches such as eddy in geometric distortion correction. Video S1 displays sagittal slices of the ex vivo brain 1 data set acquired at b = 10,000 s/mm^2^ and reconstructed using dynamic field monitoring and 1D LPC with and without geometric correction. Although eddy reduces geometric distortions, residual misalignment is still be visible in the frontal lobe and the region below the hippocampus, as indicated by the red arrows in Video S1. Residual distortions can also be seen in the enlarged sagittal image of the hippocampus (Video S2) and the temporal lobe of ex vivo brain 2 dataset (Video S3). Diffusion-weighted images reconstructed using dynamic field monitoring exhibit superior spatial alignment with no discernible residual distortions.

It is important to note that image resampling in the ‘eddy’ tool introduces blurring in the corrected images, which is noticeable when comparing ‘eddy’-corrected images with those reconstructed with 1D LPC and no additional post-processing (Figure 6A). Spatial alignment between diffusion-weighted images is required prior to voxel-wise dMRI model fitting. Dynamic field monitoring-based image reconstruction achieves accurate spatial alignment between diffusion-weighted images through eliminating geometric distortions and without incurring blurring in the resulting images. Note the more precise delineation of the claustrum and sharper contrast in the cortex (Figure 6B, yellow and green arrows, respectively) in the images reconstructed using the measured field trajectories as compared to the 1D LPC images with and without ‘eddy’ applied. Quantification of the degree of image blur with the CPBD and PSI metrics confirmed the visual observations (Figure 6C). The CPBD and PSI were consistently lower when images were reconstructed with 1D LPC and processed with ‘eddy’ than when images were reconstructed using 1D LPC without ‘eddy’ applied. The CPBD and PSI were the highest for images reconstructed with dynamic field monitoring.

**Figure 6.**
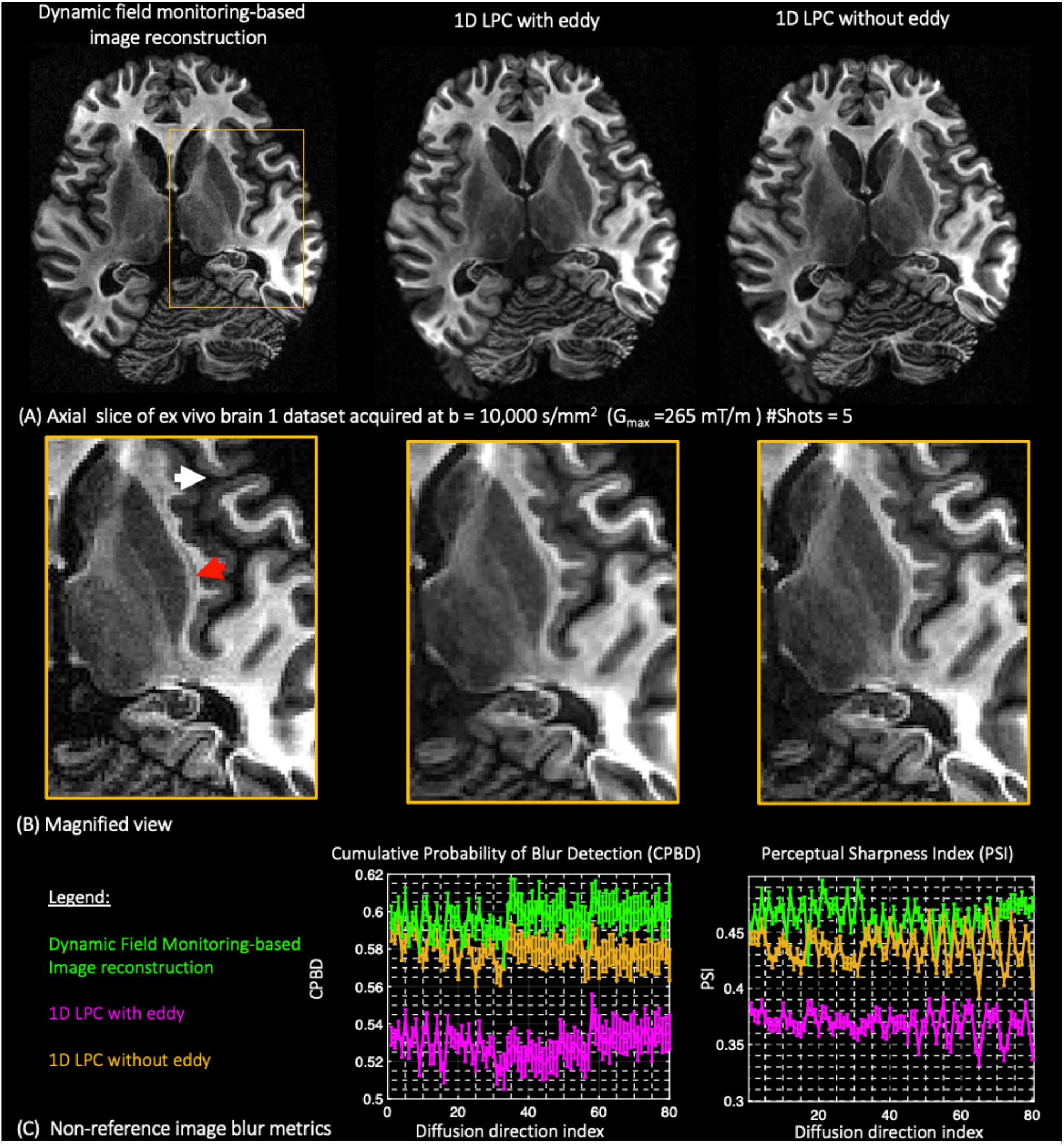
Image sharpness in diffusion-weighted images (A-B) when geometric distortions are accounted for with dynamic field monitoring reconstruction and applying ‘eddy’ after 1D LPC. For comparison, images reconstructed using 1D LPC without applying ‘eddy’ are also shown. (C) Image blur metrics confirm that eddy introduces additional blurring in contrast to dynamic field monitoring-based image reconstruction. Note the enhanced contrast in the claustrum (red arrow) and cortex (white arrow) when images are reconstructed with dynamic field monitoring.

Figure 7 shows MD, FA, and color-encoded FA maps of the ex vivo brain 1 dataset reconstructed with dynamic field monitoring-based image reconstruction, 1D LPC, and MUSE. Ghosting artifacts were pronounced in the MD maps (Figure 7A) derived from the 1D LPC and MUSE images but not observed win the images reconstructed with dynamic field monitoring. Replicas of brain structures were clearly seen in the PLP background (the hyperintense region surrounding the brain) and in the gray and white matter. Note the artifactual areas of increased MD in the brainstem of the images reconstructed with 1D LPC and MUSE (green arrow), as well as the artifacts in the cortex (orange arrow), which are not present on the images reconstructed using dynamic field monitoring. The tracts in the right cerebral hemisphere were poorly defined in the FA maps obtained after 1D LPC and MUSE reconstruction and appear clearer in FA maps from dynamic field monitoring (red arrow in Figure 7B). Marked ghosting artifacts in the cerebellum (Figure 4A) significantly influence the primary diffusion tensor eigenvector information. The color-encoded FA map for 1D LPC reveals an abnormal primary diffusion orientation pattern in the right side of the cerebellum, and fine white matter cerebellar tracts outlined in the color-encoded FA map of dynamic field monitoring (white arrow in Figure 7C) are almost indistinguishable with 1D LPC. The DTI maps from the ex vivo brain 2 datasets yield similar results (Figures S7).

**Figure 7.**
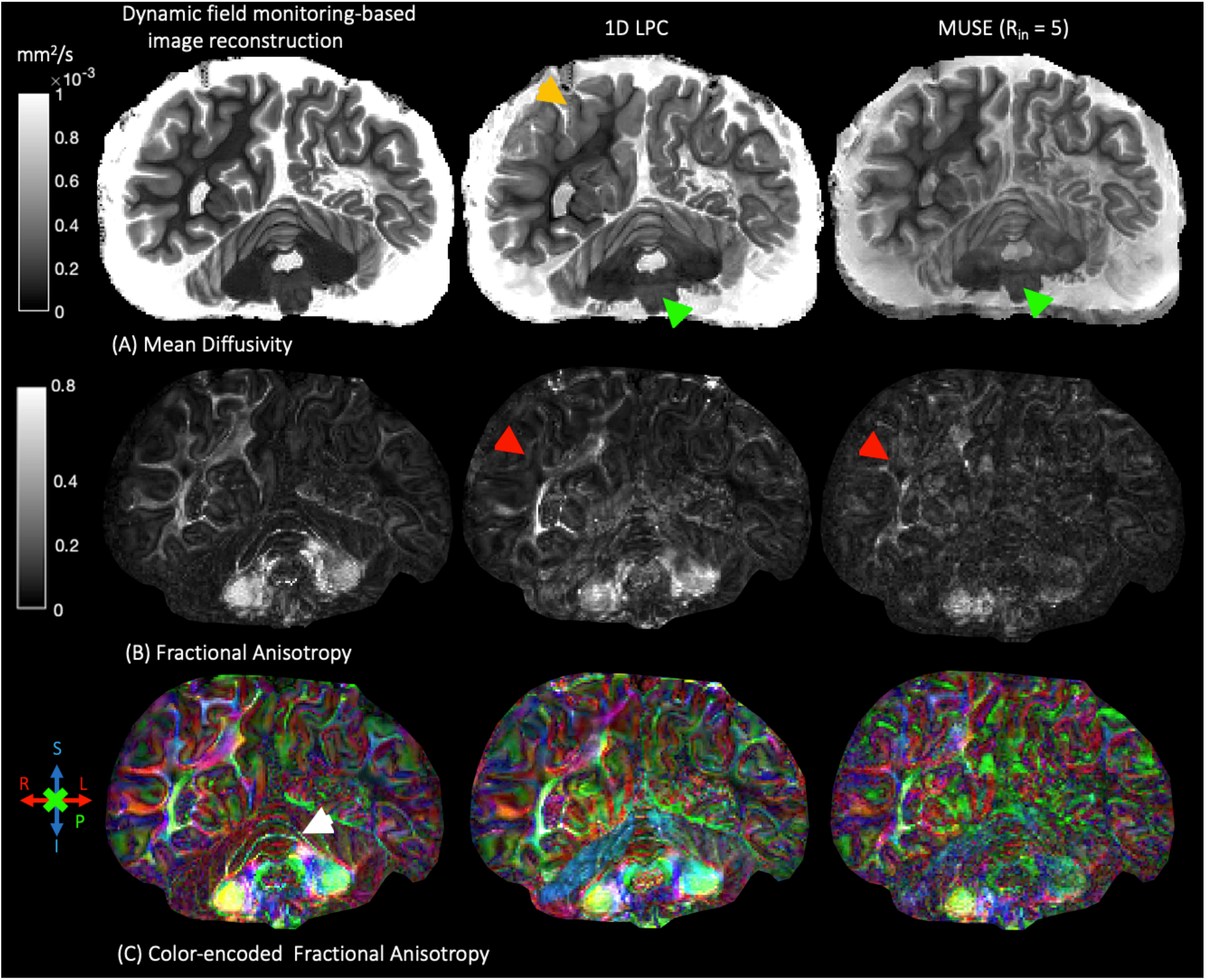
Mean diffusivity, fractional anisotropy, and color-encoded fractional anisotropy maps obtained after reconstructing diffusion-weighted images of the axial slice in Figure 4A (ex vivo brain 1 dataset) for dynamic field monitoring-based reconstruction, 1D LPC, and MUSE. Note the artifacts in the cortex (yellow arrow) and in the brainstem (green arrow) of the MD maps obtained with 1D LPC and MUSE. Red arrows point to missing tracts in FA maps from these two methods. Note the finer delineation of cerebellum fibers (white arrow in (C)) with dynamic field monitoring-based reconstruction.

Table 2 reports the mean MD and FA measured in three representative white matter tract ROIs in the corpus callosum, corona radiata, and temporal lobe. Compared to 1D LPC, we noticed a consistent increase in FA and decrease in MD when we reconstructed the images using dynamic field monitoring-based image reconstruction. The findings are in agreement with trends observed in DTI metrics calculated before and after ghosting reduction in high-gradient strength ex vivo dMRI images reconstructed using the SLM-based reconstruction method.^35^

**Table 2.**
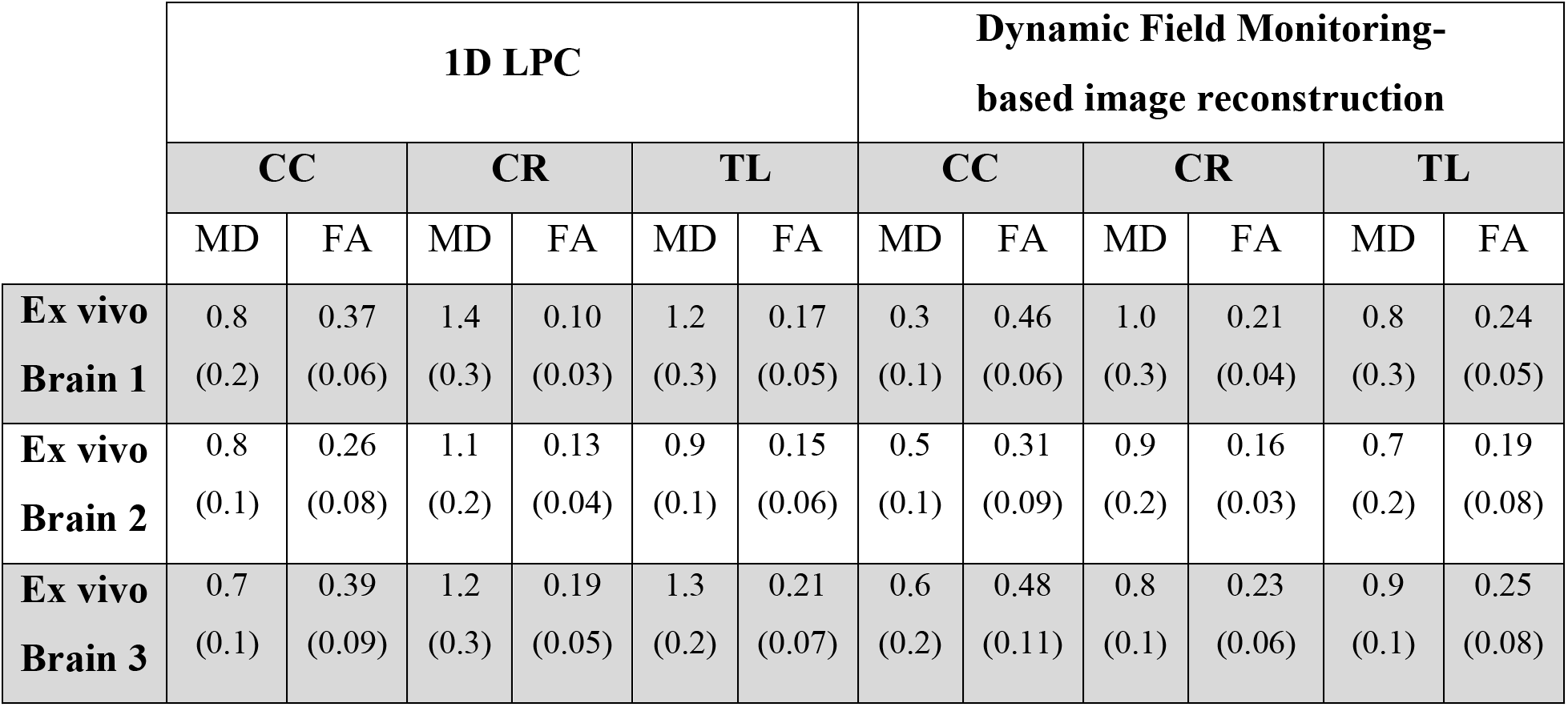
Estimated MD (mm^2^ /s x 10^-4^) and FA values in three representative white matter tract regions of interest: the corpus callosum (CC), corona radiata (CR),and the left temporal lobe (TL), after dMRI image reconstruction with the different techniques. Value in parenthesis represents standard deviation.

Figure 8 shows fiber orientation distribution functions (ODFs) calculated from the multi-shell data in the ex vivo brain 1 dataset alongside the color-encoded FA maps of the coronal slice shown in Figure 4B. The color-encoded FA maps reveal significant variation in diffusion orientation throughout the gray and white matter, with a disproportionately large number of voxels revealing primary diffusion orientation along the slab direction (A>P, green color) and only sparingly along the frequency encoding direction (R>L, red color). This is especially noticeable in the corpus callosum (Figure 8B) and the left temporal lobe (Figure 8C). Another visual observation consistent with our previous work,^35^ is that when ghosting artifacts are not entirely removed, the amplitudes of the fiber ODFs are consistently lower (compare fiber ODFS in corpus callosum of the 1D LPC maps with those of dynamic field monitoring).

**Figure 8.**
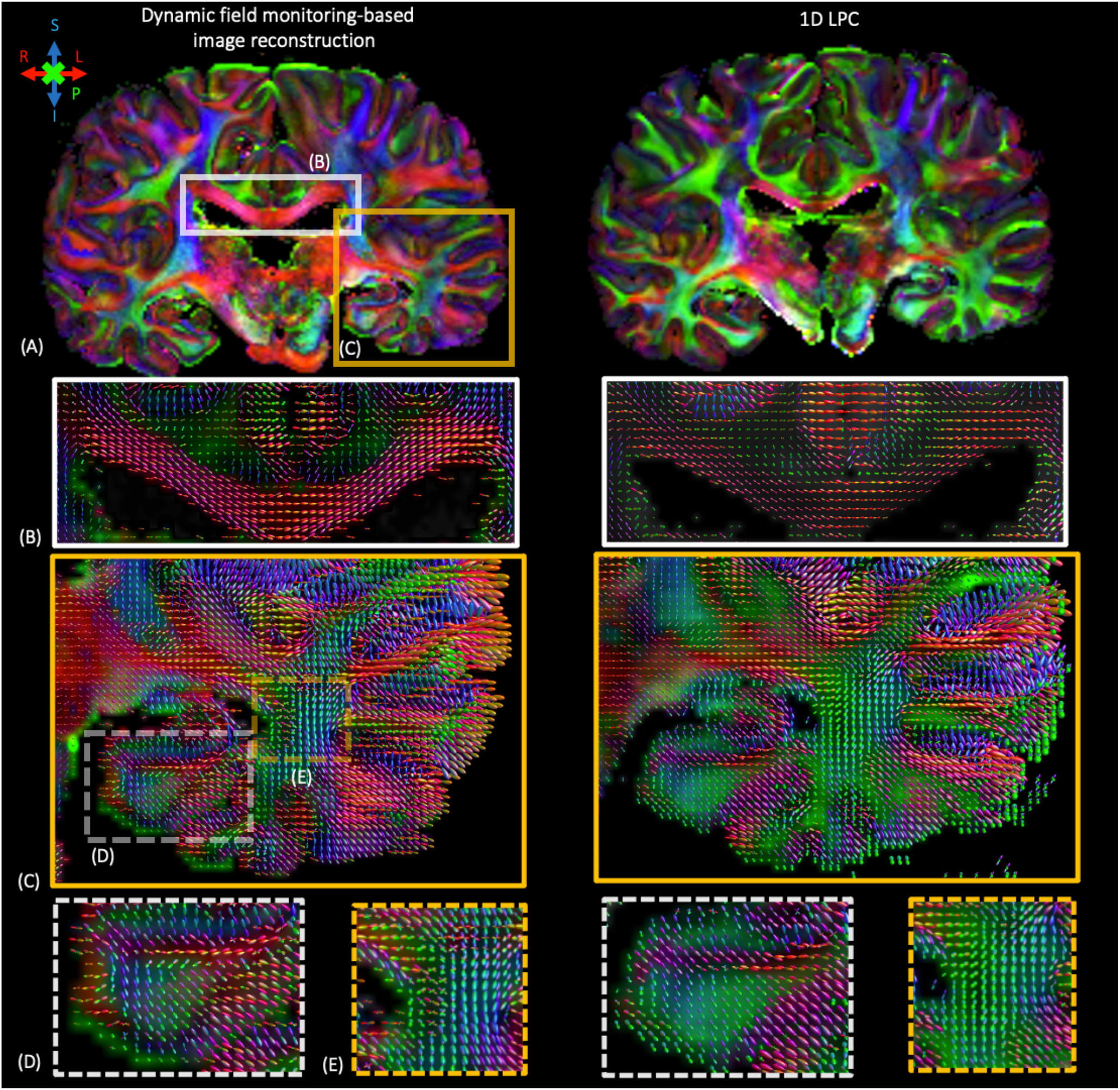
(A) Color-encoded FA maps for the same coronal slice shown in Figure 4B for ex vivo dataset 1. (B-E) Fiber ODFs in the corpus callosum (B) and temporal lobe (C) obtained after reconstructing images with dynamic field monitoring and 1D LPC. Magnified areas in the parahippocampal region (D) and in the temporal lobe (E) showing two-fiber ODFs are better visualized on the images reconstructed with dynamic field monitoring compared to LPC.

Furthermore, ghosting artifacts and insufficient geometric distortions correction in areas around the hippocampus (see Video S1 and S2) cause a loss of detail in the fiber ODFs maps of the parahippocampal region (Figure 8C). While the fiber ODFs produced after dynamic field monitoring varied in their principal orientation with respect to the depth of the entorhinal cortex (Figure 8D), such spatial topographies appeared to be missing in the fiber ODFs obtained from images reconstructed with 1D LPC, which show a substantial proportion of fiber ODFs pointing in the A>P direction. The number of crossing fibers detected also increased when dynamic field monitoring was used to reconstruct the images. Figure 8E shows an enlarged region where two-fiber ODFs can be resolved after dynamic field monitoring-based reconstruction but not after 1D LPC. The percentage of complex fiber configurations (> 1 peak) in the ex vivo brain 1 dataset’s white matter was 65% in contrast to 55% when 1D LPC and eddy registration were used. Similar values were found in the ex vivo brain 3 dataset: 68% and 55%.

## 4. Discussion

In this work, we have shown that the eddy currents introduced by using diffusion gradients up to 300 mT/m on a 3T Connectom scanner are highly nonlinear in space and time. They may extend into the image readout period, thereby interfering with the expected trajectory of the imaging gradients during the readout, as demonstrated here in a 3D diffusion-weighted multi-shot EPI sequence. Eddy current fields introduce phase modulations between odd and even EPI echoes and between shots that are of high-spatial order besides changing dynamically during the readout. Dynamic field monitoring-based image reconstruction is an effective way to correct ghosting and geometric distortion artifacts incurred by such eddy currents. By measuring the nonlinear phase evolution during the readout period and incorporating this information into a multi-shot image reconstruction forward model, we successfully reconstructed submillimeter resolution (~0.8 mm isotropic) diffusion-weighted images using b-values up to 10,000 s/mm^2^ with more effective reduction of ghosting than current state-of-the-art reconstruction approaches, and with more accurate geometric distortion correction than popular image-based approaches such as ‘eddy’. Both ghosting and geometric distortion artifacts were handled simultaneously in a single step, avoiding artifact propagation in the processing pipeline. Images reconstructed with our method yield DTI and fiber ODFs vector maps that are free of orientational bias.

We chose to demonstrate the benefits of dynamic field monitoring in high-gradient dMRI of ex vivo human brain samples scanned on the Connectom scanner. The Connectom scanner and other human MRI systems featuring high-performance gradient technology^74–76^ serve as ideal platforms for performing high b-value ex vivo dMRI of whole human brain specimens, which are larger than can be accommodated on preclinical imaging systems. Diffusion imaging of large ex vivo tissue specimens benefits from highly segmented acquisitions to achieve high spatial resolution within short echo times while utilizing strong diffusion-sensitizing gradients to achieve sufficient diffusion contrast. This data, paired with ground truth from microscopy, will empower the development of new tools for inferring axonal orientations and microstructure from dMRI.^41,77^

As gradient performance and technology improve and become integrated into commercially available human MRI systems^78^, we anticipate that the results of this paper will inform future efforts to maximize the use of strong gradients for dMRI and other applications. ^14^ We expect that dynamic field monitoring will be an integral step to handling eddy current artifacts at the unprecedented gradient strengths levels of the new Connectome 2.0 scanner, nearly doubling the maximum gradient strength (500 mT/m vs. 300 mT/m).^14^

Previously, we showed that an extension of the structured low-rank reconstruction method^36^ to a multi-shot acquisition^35^ was effective in reducing ghosting artifacts compared to other state-of-the-art multi-shot approaches at a high-gradient strength regime.^34^ The structured low-rank reconstruction method and other approaches such as MUSE underperform in ghost correction when the number of shots was high (>5). Such non-linear ghost correction approaches rely on sufficient encoding power (i.e., number of coil channels) to solve an undersampled reconstruction problem and make use of regularization and prior information to compensate for the lack of k-space measurements.^34,79–83^ When the number of shots and the spatial resolution is high, the reconstruction problem cannot be solved accurately. Another simplistic assumption in such models is that a static phase map (possibly non-linear in space) can characterize either the set of odd or even k-space lines or for each shot ^84,85^. However, as we have shown here, the phase modulation between odd and even k-space lines may not be constant during the readout, with potential for large dynamic variations in the magnetic field due to the presence of residual eddy current fields.

As for geometric distortions, we demonstrated that more accurate spatial alignment across diffusion-weighted volumes could be obtained if dynamic field monitoring is used instead of post-processing software tools like eddy. Even if satisfactory geometric distortion correction is achieved with eddy, image resampling in the registration step will introduce image blurring. Sharper images can be obtained with dynamic field monitoring reconstruction. While the degree of image blurring may be discounted for low-spatial resolution acquisitions, the effect of blurring becomes increasingly important at high spatial resolution, as demonstrated here in the submillimeter dMRI acquisitions targeted at evaluating fine white matter tracts and the cortex,^86–89^ where partial volume effects due to blurring can confound dMRI analysis in such structures.

In this work, we assumed that phase variations along partitions arise solely from the spherical harmonic coefficient associated with *h*_3_ (***r***) = *z*. Such variations are attributed to *k_z_* phase increments of the nominal trajectory and possibly linear eddy currents along *z*. The remaining spherical harmonics coefficients are assumed to have a non-negligible variation over partitions. This assumption has been applied in our previous work satisfactorily^35^, and makes it possible to break up the reconstruction of the 3D volume into a series of 2D multi-shot reconstruction problems by applying a non-uniform Fourier transform along the slab encoding direction,^62^ rendering the reconstruction more computationally tractable. While we have obtained excellent results in the postmortem datasets shown here, we acknowledge that uncorrected ghosting and geometric artifacts may still be present. Reconstruction of the full 3D volume was not possible with our current implementation due to the high RAM requirement needed to allocate the full 3D image encoding matrix in MATLAB. For future work, we are considering including approximation methods that use singular value decomposition to reduce the dimensionality of the forward encoding matrix^90^, or on-the-fly implementation of matrix multiplication (without explicitly saving the encoding matrix in memory) in faster programming languages than MATLAB, e.g., C++ or Julia.

The choice of the Tikhonov regularization parameter λ can affect the apparent SNR of the reconstructed images. The images reconstructed with dynamic field monitoring were slightly noisier than those obtained with 1D LPC. A plausible explanation is that the image encoding matrix used in the field monitoring reconstruction had less favorable properties than the 3D Fourier matrix used in 1D LPC in terms of noise amplification. In this sense, tuning the Tikhonov regularization parameter with classical approaches as the L-curve^91^ may help in reducing the noise further, although we note that the current choice of λ did not affect the degree of ghosting and distortion correction.

The multi-shot reconstruction framework presented here preserves the Gaussian noise statistics of the raw k-space data as the reconstructed image is the output of a linear operator.^92^ De-noising algorithms that rely on Gaussian noise statistics can be applied after image reconstruction to compensate for the amplification of noise in the reconstruction if the choice of the Tikhonov regularization parameter was deemed insufficient.^93–95^

## 5. Conclusions

Eddy current-induced fields from the strong diffusion gradients applied in a 3T Connectom scanner are highly nonlinear in space and time. These eddy currents can create pronounced ghosting artifacts and geometric distortions in the resulting images, particularly for segmented EPI acquisitions with multiple shots. The artifacts are not fully addressed with the currently available image reconstruction and distortion correction methods. Integrating the actual phase evolution measured with dynamic field monitoring into the image reconstruction framework achieves ghosting, distortion and blur-free images by precisely accounting for the higher-order eddy current fields. This physics-informed approach to correcting eddy current artifacts has been validated qualitatively and quantitatively in high spatial resolution ex vivo whole human brain dMRI utilizing high b-values up to 10,000 s/mm^2^. We anticipate this approach will gain increasing importance and relevance with the adoption of high-performance gradient technology for dMRI and other applications in commercially available and cutting-edge research scanners.

## Acknowledgments

This work was supported by the National Institutes of Health (grant numbers P41-EB030006, U01-EB026996, R01-NS118187, U54NS115322), the United States Department of Defense (W81XWH2210999) and the Athinoula A. Martinos Center for Biomedical Imaging. We would like to thank Drs. Paul Weavers, Cameron Cushing, and Christian Mirkes for invaluable assistance and support during this project, and Dr. Bruce Fischl and Jocelyn Mora for sharing two of the post-mortem specimens.

## Figure captions

**Figure S1.** Position of the post-mortem brain in the 48 channel whole-brain receive array coil designed for mesopic ex vivo dMRI. Adapted from ^35^.

**Figure S2.** Zero and first-order spherical harmonic terms describing the spatiotemporal phase behavior of eddy current fields induced by diffusion gradients at G_max_ = 94 mT/m, G_max_ = 171 mT/m, G_max_ = 232 mT/m and G_max_= 277 mT/m, for the first shot and kz close to the zero frequency. The maximum phase accrual achieved within the FOV is shown in radians *ϕ_eddy-h_l__*(*t*) for every harmonic *h_l_*, (***r***). Zero and First order terms are shown.

**Figure S3.** Second-order spherical harmonic terms describing the spatiotemporal phase behavior of eddy current fields induced by diffusion gradients at G_max_ = 94 mT/m, G_max_ = 171 mT/m, G_max_ = 232 mT/m and G_max_= 277 mT/m, for the first shot and *k_z_* close to the zero frequency. The maximum phase accrual achieved within the FOV is shown in radians *ϕ_eddy-h_l__*(*t*) for every harmonic *h_l_*, (***r***). Zero and First order terms are shown.

**Figure S4.**Phase variations of eddy current fields induced by diffusion gradients at G_max_ = 277 mT/m between the third and the second shot.

**Figure S5.** Coronal slice from the ex vivo brain 2 dataset reconstructed with dynamic field monitoring-based image reconstruction, 1D LPC, MUSE, and SLM-based ghosting method at G_max_ = 200 mT/m. Neither MUSE nor SLM-method could provide acceptable images in this seven-shot EPI acquisition.

**Figure S6.** Coronal slice from the ex vivo brain 3 dataset reconstructed with dynamic field monitoring-based image reconstruction, 1D LPC, MUSE, and SLM-based ghosting method at (A) b = 4,000 s/mm^2^, G_max_ = 144 mT/m, and (B) b= 10,000 s/mm^2^, G_max_ = 268 mT/m.

**Figure S7.** Mean Diffusivity, Fractional Anisotropy, and color-encoded Fractional Anisotropy maps obtained after reconstructing diffusion-weighted images of the coronal slice in Figure S6 (ex vivo brain 2 dataset) for dynamic field monitoring-based reconstruction and 1D LPC.

## Table captions

**Table S1.** Real-Valued Solid Harmonics up to third order

## Video captions

**Video S1.** Geometric distortions artifacts correction with dynamic field monitoring and eddy for the ex vivo brain 1 dataset. A sagittal view of diffusion-weighted images at b = 10,000 s/mm^2^ and G_max_ = 270 mT/m is shown.

**Video S2.** Sagittal view of the hippocampus for the ex vivo brain 1 dataset at b = 10,000 s/mm^2^ and G_max_ = 270 mT/m where geometric distortion artifacts can still be noticed after eddy but removed with dynamic field monitoring.

**Video S3.**Geometric distortions artifacts correction with dynamic field monitoring and eddy for the ex vivo brain 2 dataset. A coronal view of diffusion-weighted images at b = 4,000 s/mm^2^ and G_max_ = 200 mT/m is shown.

