## Supplementary file for "Eddy current-induced artifacts correction in high gradient strength diffusion MRI with dynamic field monitoring: demonstration in ex vivo human brain imaging"

2.1 **Nonlinear phase modeling during image encoding**

Table S1 Real-Valued Solid Harmonics up to third order

| Solid Spherical Harmonics | Order |
| --- | --- |
| $\boldsymbol{h}_{\boldsymbol{0}}\left( \boldsymbol{r} \right)\boldsymbol{=1}$ | 0 |
| $\boldsymbol{h}_{\boldsymbol{1}}\left( \boldsymbol{r} \right)\boldsymbol{=x}$  $\boldsymbol{h}_{\boldsymbol{2}}\left( \boldsymbol{r} \right)\boldsymbol{=y}$  $\boldsymbol{h}_{\boldsymbol{3}}\left( \boldsymbol{r} \right)\boldsymbol{=z}$ | 1 |
| $\boldsymbol{h}_{\boldsymbol{4}}\left( \boldsymbol{r} \right)\boldsymbol{=xy}$  $\boldsymbol{h}_{\boldsymbol{5}}\left( \boldsymbol{r} \right)\boldsymbol{=zy}$  $\boldsymbol{h}_{\boldsymbol{6}}\left( \boldsymbol{r} \right)\boldsymbol{=3}\boldsymbol{z}^{\boldsymbol{2}}\boldsymbol{-(}\boldsymbol{x}^{\boldsymbol{2}}\boldsymbol{+}\boldsymbol{y}^{\boldsymbol{2}}\boldsymbol{+}\boldsymbol{z}^{\boldsymbol{2}}\boldsymbol{)}$  $\boldsymbol{h}_{\boldsymbol{7}}\left( \boldsymbol{r} \right)\boldsymbol{=xz}$  $\boldsymbol{h}_{\boldsymbol{8}}\left( \boldsymbol{r} \right)\boldsymbol{=}\boldsymbol{x}^{\boldsymbol{2}}\boldsymbol{-}\boldsymbol{y}^{\boldsymbol{2}}$ | 2 |
| $\boldsymbol{h}_{\boldsymbol{9}}\left( \boldsymbol{r} \right)\boldsymbol{=3}\boldsymbol{y}\boldsymbol{x}^{\boldsymbol{2}}\boldsymbol{-}\boldsymbol{y}^{\boldsymbol{3}}$  $\boldsymbol{h}_{\boldsymbol{10}}\left( \boldsymbol{r} \right)\boldsymbol{=xyz}$  $\boldsymbol{h}_{\boldsymbol{11}}\left( \boldsymbol{r} \right)\boldsymbol{=}\left( \boldsymbol{5}\boldsymbol{z}^{\boldsymbol{2}}\boldsymbol{-}\left( \boldsymbol{x}^{\boldsymbol{2}}\boldsymbol{+}\boldsymbol{y}^{\boldsymbol{2}}\boldsymbol{+}\boldsymbol{z}^{\boldsymbol{2}} \right) \right)\boldsymbol{y}$  $\boldsymbol{h}_{\boldsymbol{12}}\left( \boldsymbol{r} \right)\boldsymbol{=5}\boldsymbol{z}^{\boldsymbol{3}}\boldsymbol{-3}\boldsymbol{z}\left( \boldsymbol{x}^{\boldsymbol{2}}\boldsymbol{+}\boldsymbol{y}^{\boldsymbol{2}}\boldsymbol{+}\boldsymbol{z}^{\boldsymbol{2}} \right)$  $\boldsymbol{h}_{\boldsymbol{13}}\left( \boldsymbol{r} \right)\boldsymbol{=}\left( \boldsymbol{5}\boldsymbol{z}^{\boldsymbol{2}}\boldsymbol{-}\left( \boldsymbol{x}^{\boldsymbol{2}}\boldsymbol{+}\boldsymbol{y}^{\boldsymbol{2}}\boldsymbol{+}\boldsymbol{z}^{\boldsymbol{2}} \right) \right)\boldsymbol{x}$  $\boldsymbol{h}_{\boldsymbol{14}}\left( \boldsymbol{r} \right)\boldsymbol{=}\boldsymbol{x}^{\boldsymbol{2}}\boldsymbol{z-}\boldsymbol{y}^{\boldsymbol{2}}\boldsymbol{z}$  $\boldsymbol{h}_{\boldsymbol{15}}\left( \boldsymbol{r} \right)\boldsymbol{=}\boldsymbol{x}^{\boldsymbol{3}}\boldsymbol{-3}\boldsymbol{x}\boldsymbol{y}^{\boldsymbol{2}}$ | 3 |

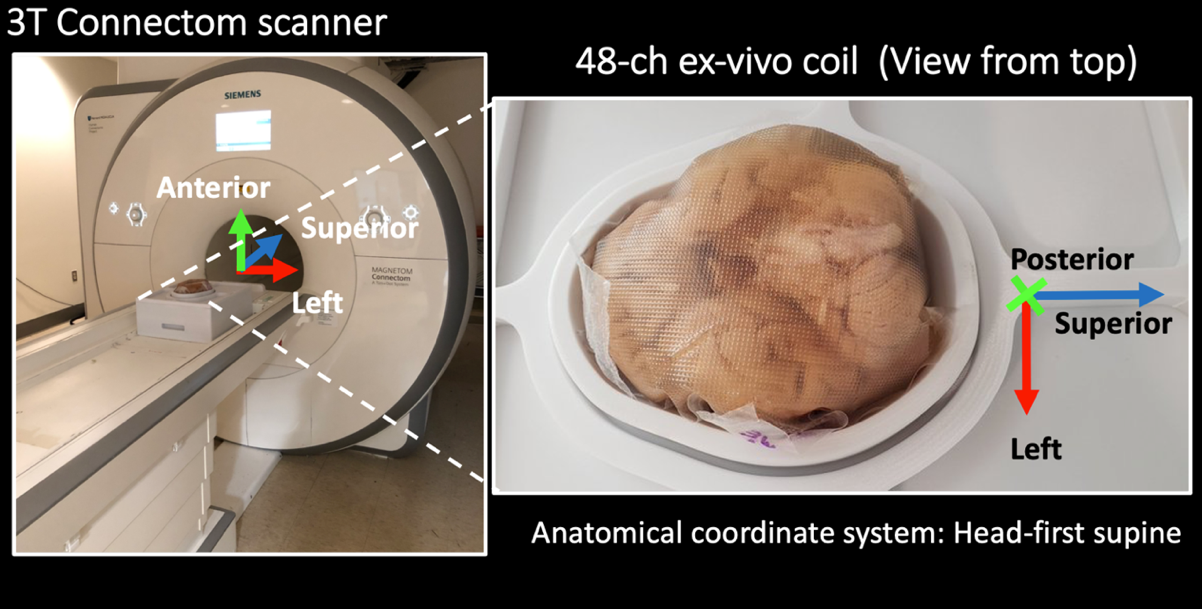
**2.4.2 Experiment 2: Correction of ghosting and geometric distortions induced by eddy currents in high gradient strength dMRI experiments on ex vivo human brain samples**

Figure S1 Position of the post-mortem brain in the 48 channel whole-brain receive array coil designed for mesopic ex vivo diffusion MRI. Adapted from ^1^.

The head-first supine anatomical coordinate system (inferior–superior, left–right, anterior–posterior) in relation to the ex vivo brain is indicated in the figure. Note that with this convention, the neuroanatomical coordinates of the brain are related to the anatomical coordinate system as follows: ventral–dorsal is anterior–posterior, left–right is right–left, and rostral–caudal is inferior–superior.

**3.1 Experiment 1: Characterization of eddy current fields induced by diffusion-sensitizing gradients**

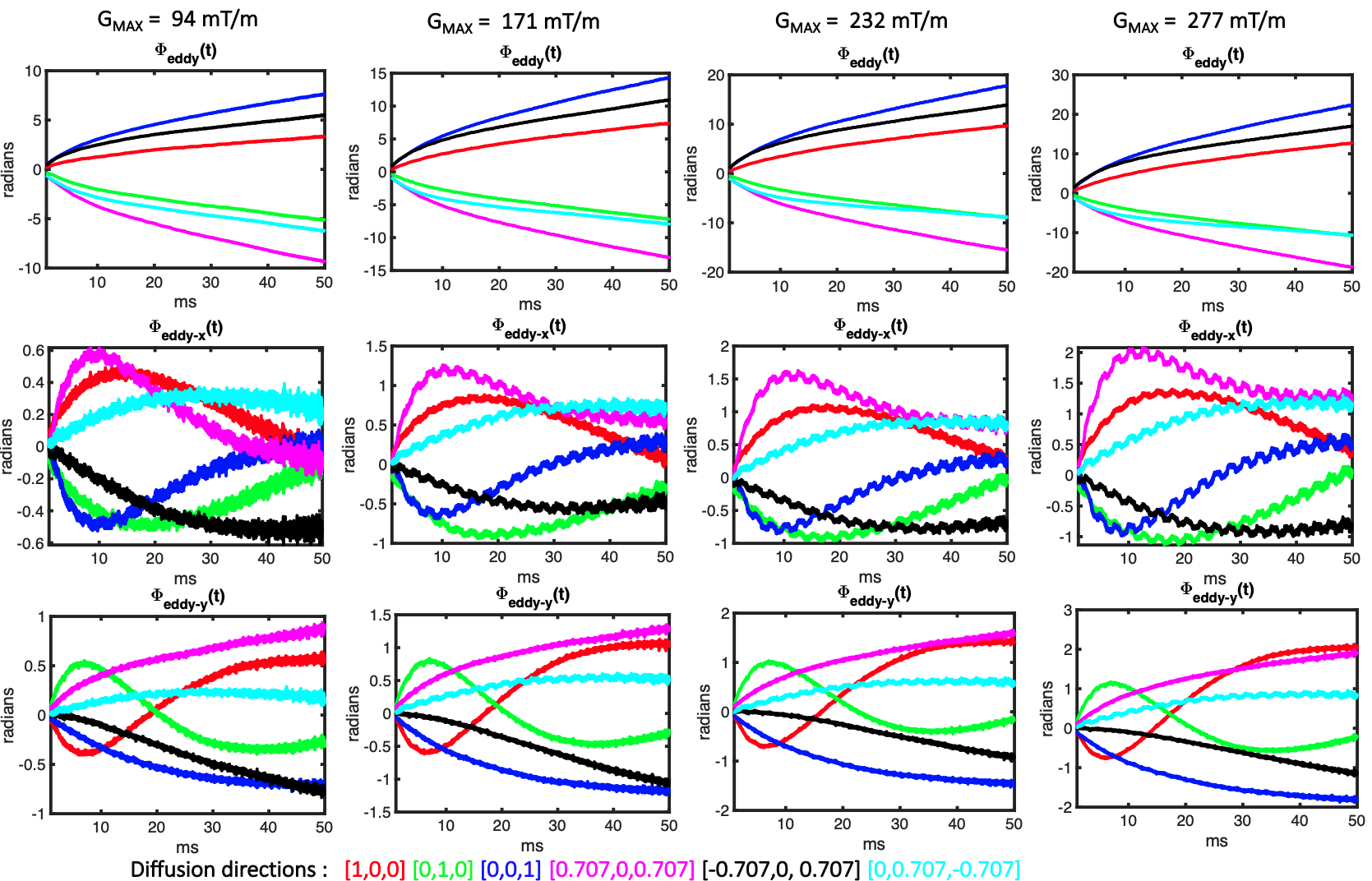

Figure S2 Zero and first-order spherical harmonic terms describing the spatiotemporal phase behavior of eddy current fields induced by diffusion gradients at G_max_ = 94 mT/m, G_max_ = 171 mT/m, G_max_ = 232 mT/m and G_max_= 277 mT/m, for the first shot and k_z_ close to the zero frequency. The maximum phase accrual achieved within the FOV is shown in radians $\phi_{eddy-h_{l}}\left( t \right)$ for every harmonic $h_{l}(\mathbf{r})$. Zero and First order terms are shown.

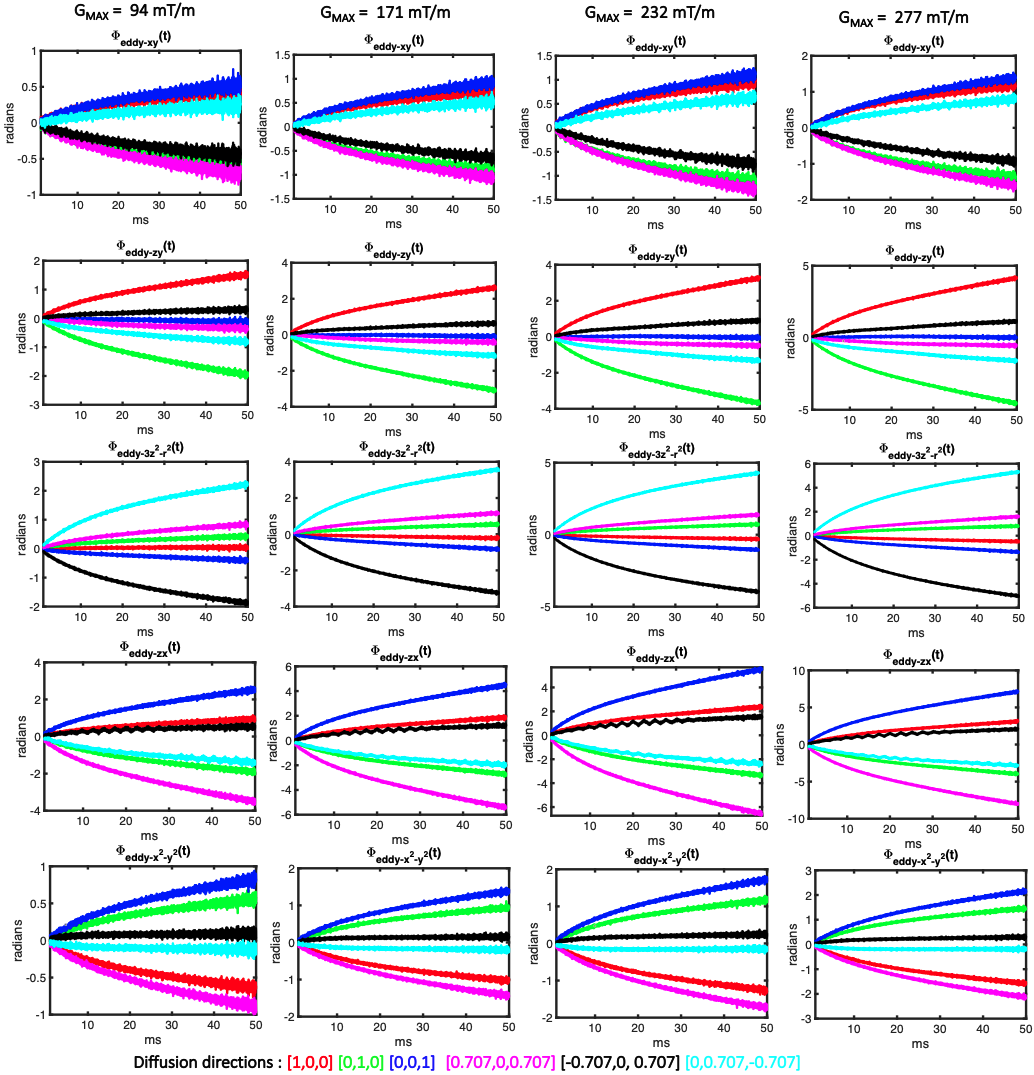

Figure S3 Second order spherical harmonic terms describing the spatiotemporal phase behavior of eddy current fields induced by diffusion gradients at G_max_ = 94 mT/m, G_max_ = 171 mT/m, G_max_ = 232 mT/m and G_max_= 277 mT/m, for the first shot and kz close to the zero frequency. The maximum phase accrual achieved within the FOV is shown in radians $\phi_{eddy-h_{l}}\left( t \right)$ for every harmonic $h_{l}(\boldsymbol{r})$. Zero and First order terms are shown.

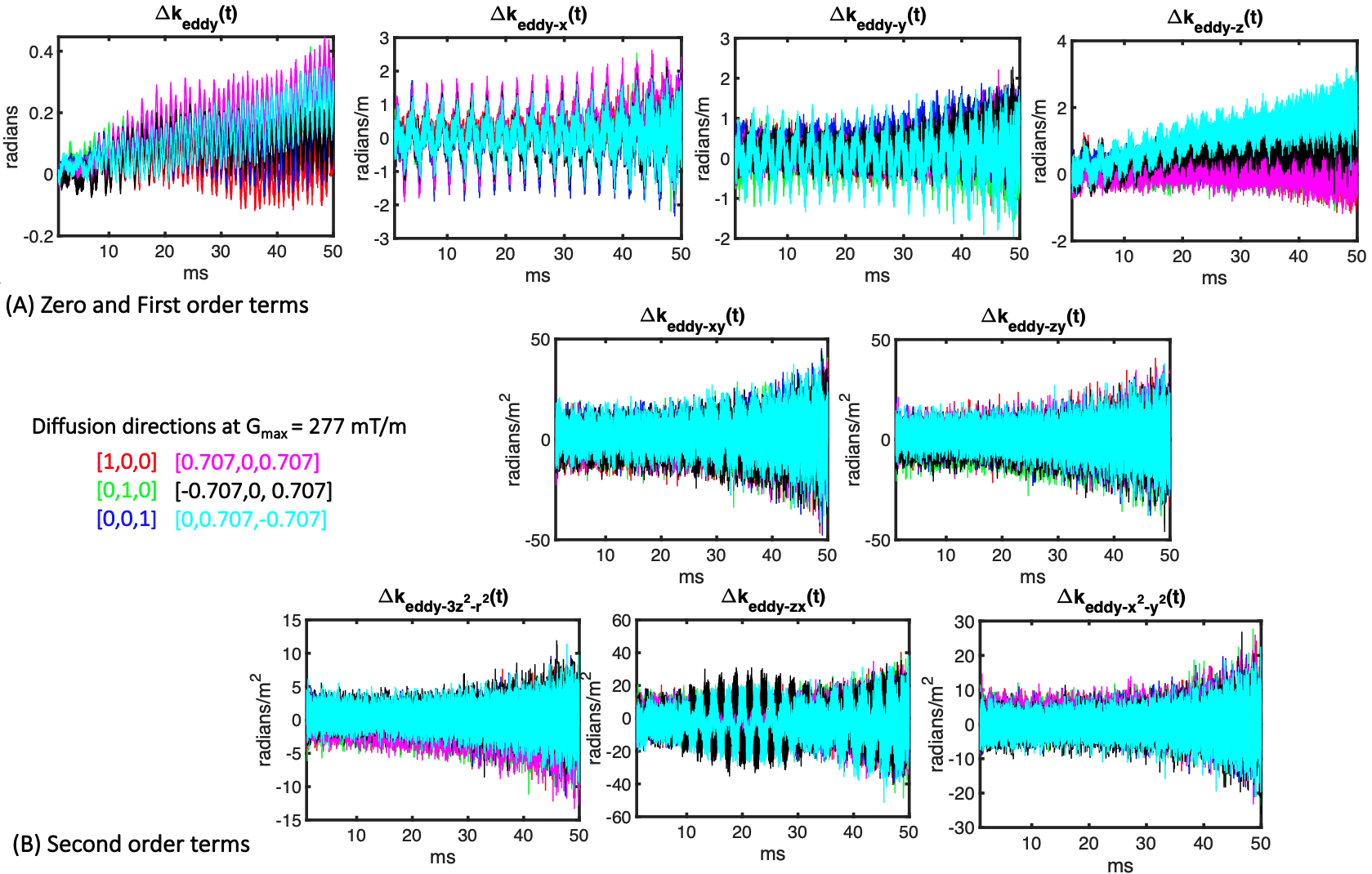

Figure S4 Phase variations of eddy current fields induced by diffusion gradients at G_max_ = 277 mT/m between the third and the second shot.

**3.2 Experiment 2: Correction of ghosting and geometric distortions induced by eddy currents in high gradient strength dMRI experiments on ex-vivo human brain samples**

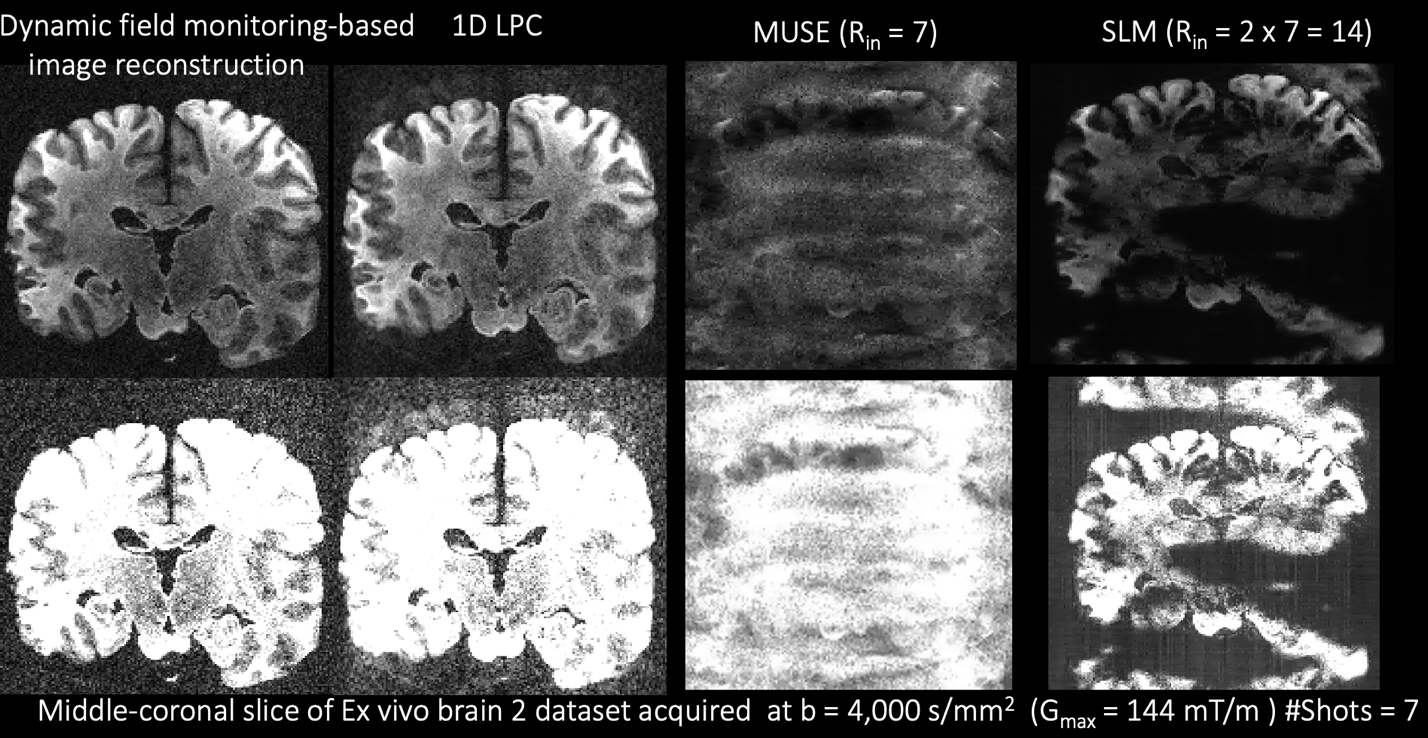

Figure S5 Coronal slice from the ex vivo brain 2 dataset reconstructed with dynamic field monitoring-based image reconstruction, 1D LPC, MUSE, and SLM-based ghosting method at G_max_ = 200 mT/m. Neither MUSE nor SLM-method could provide acceptable images in this seven-shot EPI acquisition.

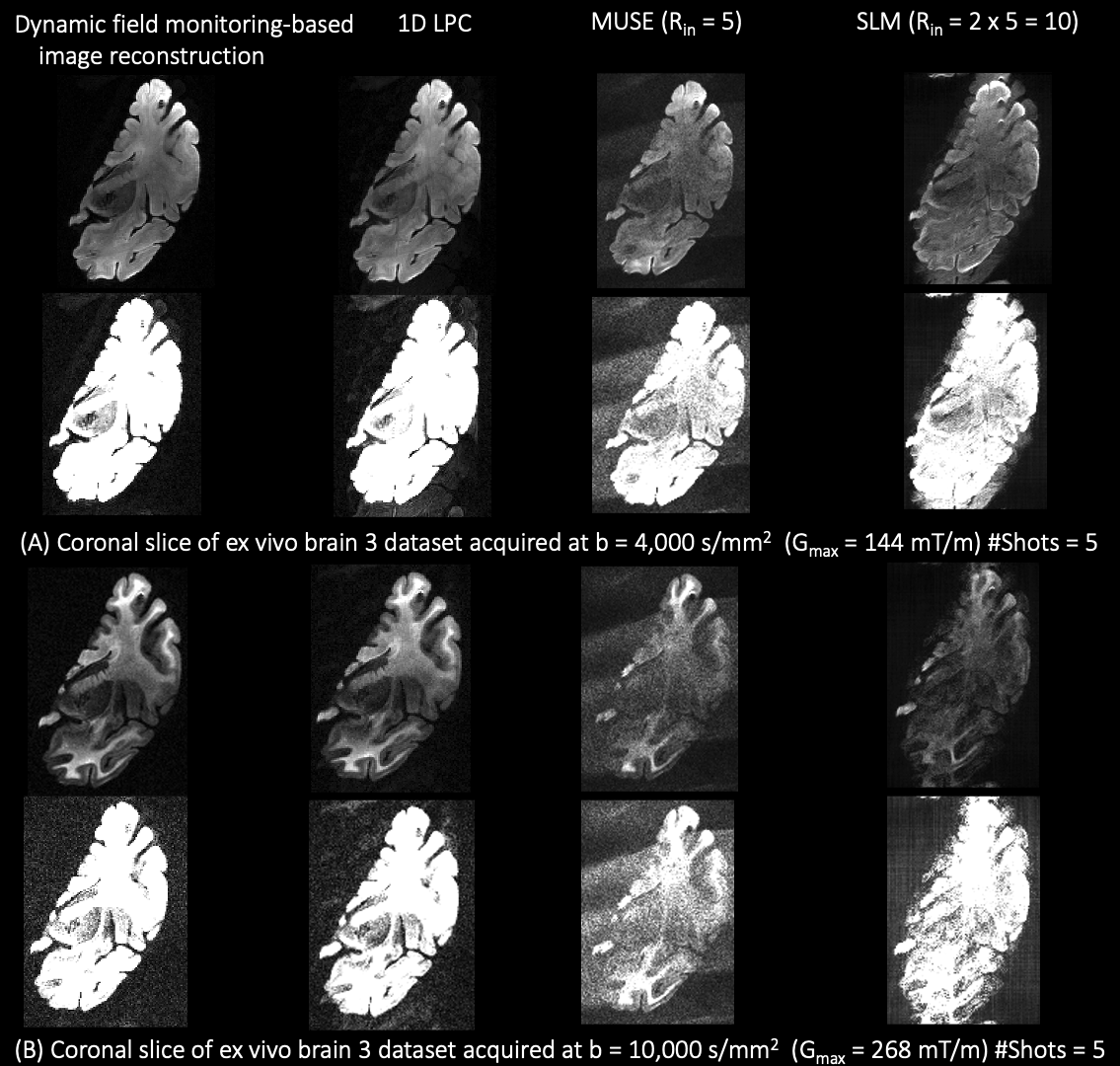

Figure S6. Coronal slice from the ex vivo brain 3 dataset reconstructed with dynamic field monitoring-based image reconstruction, 1D LPC, MUSE, and SLM-based ghosting method at (A) b = 4,000 s/mm^2^ , G_max_ = 144 mT/m, and (B) b= 10,000 s/mm^2^ , G_max_ = 268 mT/m.

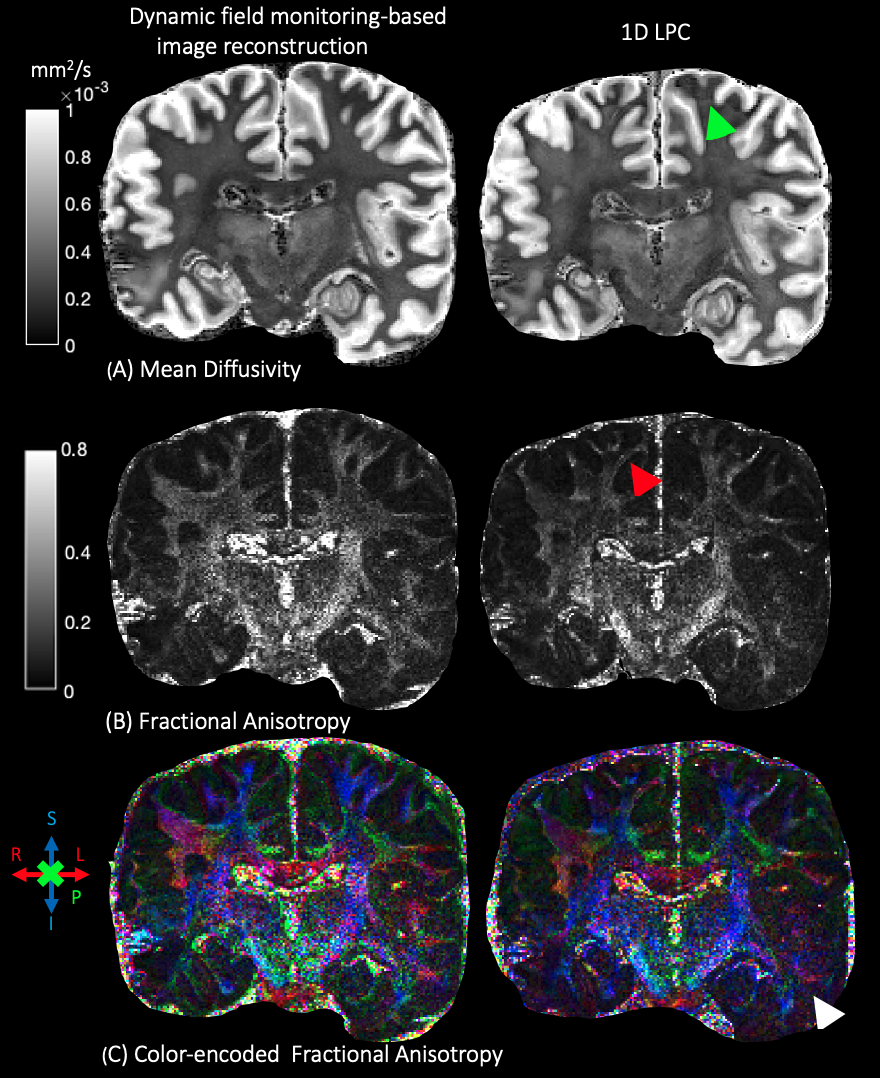

Figure S7. Mean Diffusivity, Fractional Anisotropy, and color-encoded Fractional Anisotropy maps obtained after reconstructing diffusion-weighted images of the coronal slice in Figure S6 (ex vivo brain 2 dataset) for dynamic field monitoring-based reconstruction and 1D LPC.
